# Cholesteryl hemiazelate Identified in Cardiovascular Disease Patients Causes *in vitro* and *in vivo* Inflammation

**DOI:** 10.1101/2023.02.09.527823

**Authors:** Neuza Domingues, Joana Gaifem, Rune Matthiesen, Diana P. Saraiva, Luís Bento, André R.A. Marques, Maria I. L. Soares, Julio Sampaio, Christian Klose, Michal A. Surma, Manuel S. Almeida, Gustavo Rodrigues, Pedro Araújo Gonçalves, Jorge Ferreira, Ryan Gouveia e Melo, Luís Mendes Pedro, Kai Simons, Teresa M. V. D. Pinho e Melo, M. Guadalupe Cabral, Antonio Jacinto, Ricardo Silvestre, Winchil Vaz, Otília V. Vieira

## Abstract

Oxidation of polyunsaturated fatty acids (PUFA) in low-density lipoproteins (LDL) trapped in the arterial intima plays a critical role in atherosclerosis. Though there have been many studies on the atherogenicity of oxidized derivatives of unsaturated fatty acid esters of cholesterol, the effects of the oxidation end-products of these esters has been ignored in the literature.

Through lipidomics analyses of the plasma of cardiovascular disease patients and human endarterectomy specimens we identified and quantified cholesteryl hemiesters (ChE), end-products of oxidation of polyunsaturated-fatty acid esters of cholesterol. Cholesteryl hemiazelate (ChA) was the most prevalent ChE identified. Importantly human monocytes, monocyte-derived macrophages (MDM) and neutrophils exhibit inflammatory features when exposed to sub-toxic concentrations of ChA *in vitro*. ChA increases the secretion of proinflammatory cytokines such as IL-1β and IL-6 and modulates the surface markers profile of monocytes and MDM. *In vivo*, when zebrafish larvae were fed with a ChA-enriched diet they exhibited neutrophil and macrophage accumulation in the vasculature in a caspase 1- and cathepsin B-dependent manner. ChA also triggered lipid accumulation at the bifurcation sites of the vasculature of the zebrafish larvae and negatively impacted their life expectancy.

We conclude that ChA has pro-atherogenic properties and can be considered part of a damage-associated molecular pattern (DAMP) in the development of atherosclerosis.

## INTRODUCTION

The underlying cause of most cardiovascular diseases (CVD) is atherosclerosis, a chronic progressive inflammatory pathology characterized by sub-intimal lipid accumulation in the vascular wall that can progress for years without symptoms (Libby 2002; Hansson, Libby et al. 2015). The relationship between low-density lipoprotein (LDL) cholesterol and risk of CVD has been well established through numerous epidemiological observational and genetic studies and interventional clinical trials (Libby 2002; Goldstein and Brown 2015). Since decades, oxidation of accumulated unsaturated lipids in the arterial intima has been believed to be the cause of most of the pathological consequences of atherosclerosis. Though generalised targeting of lipid peroxidation with antioxidants has not proven successful in the prevention of CVD.(McCullough, Peterson et al. 2012; Vogiatzoglou, Mulligan et al. 2015), the fact that transgenic transgenic expression of a natural antibody to oxidized phospholipids suppresses lesions in mice argues in favor of the lipid oxidation playing a critical role in atherosclerosis (Que, Hung et al. 2018). Indeed, a large body of evidence suggests that the oxidation of polyunsaturated fatty acids (PUFA) in LDL trapped in the vessel wall produce proinflammatory lipid species leading to 1) the expression of adhesion molecules and chemotactic factors that contribute to the recruitment of circulating monocytes into the intimal space, 2) the inhibition of the ability of resident macrophages to leave the intima, 3) enhancing lipoprotein uptake rate, which leads to lysosomal lipid accumulation and to “pathogenic” foam cell formation, and 4) increasing cell death and consequently loss of endothelial integrity, leukocyte recruitment and inflammation (Quinn, Parthasarathy et al. 1985; Stocker and Keaney 2004; Schmitz and Grandl 2009).

Among the oxidized LDL (oxLDL) components, oxidized phospholipids (oxPL), oxidized PUFA-esters of cholesterol (PUFA-CE) and oxysterols have been the major focus of several studies in the field, resulting in the identification of molecular structures, receptors and signaling pathways associated with inflammatory responses in vascular cells (Colles, Maxson et al. 2001; Podrez, Poliakov et al. 2002; Kadl, Sharma et al. 2011; Lee, Birukov et al. 2012; Salomon 2012; Leibundgut, Witztum et al. 2013; Rao, Zhong et al. 2014; Luu, Sharpe et al. 2016; Oh, Zhang et al. 2016). However, in all these studies, very little attention has been given to cholesteryl hemiesters (ChE), the end products of oxidation of the PUFA-CE.

LDL transport cholesteryl esters (CE), 70% of which are PUFA esters and on average, 23% of cholesteryl linoleate, 16% of cholesteryl arachidonate and 12% of cholesteryl docosahexaenoate are oxidized in human atherosclerotic lesions (Hutchins, Moore et al. 2011; Ravandi, Leibundgut et al. 2014). Oxidative scission of PUFA-CE leads to the formation of oxo-esters of cholesterol which have been identified in the “core aldehyde” fraction of *ex vivo* samples of human atheromata (Hutchins, Moore et al. 2011) and also in oxLDL (Kamido, Kuksis et al. 1995). As we have argued earlier (Estronca, Silva et al. 2012), oxo-esters of cholesterol (“core aldehydes”) can be expected to be further oxidized to the corresponding hemiesters of cholesterol. ChE are amphiphilic and have a negative charge at physiological pH. Similarly to what has been demonstrated for cholesterol (Estronca, Filipe et al. 2014), ChE may also be expected to equilibrate through passive diffusion between the oxLDL (where they are formed) and the aqueous (extra- and intracellular) phases, as well as all membranes of neighboring cells. However, despite the well estabilished presence of their percursors in the atheroma, the role of ChE in atherogenesis is still unclear. We have previously shown that a commercially available, but physiologically irrelevant ChE, cholesteryl hemisuccinate (ChS), is sufficient to cause irreversible lysosomal lipid accumulation (lipidosis) and is toxic to macrophages (Estronca, Silva et al. 2012). In primary mouse bone marrow-derived macrophages (BMDM), ChS-exposure increased the secretion of IL-1β, TNF-α and IL-6 (Domingues, Estronca et al. 2017). All these features are typically seen in atherosclerosis. Additionally, zebrafish larvae fed with a ChS-enriched diet exhibited lipid accumulation, myeloid cell-infiltration into the vasculature and decrease in life expectancy (Domingues, Estronca et al. 2017). Interestingly, under the same conditions the effects of ChS were more profound than the effects of free cholesterol (FC) (Domingues, Estronca et al. 2017). More recently, we synthesized cholesteryl hemiazelate (ChA), a cholesteryl-9-oxononanoate oxidation product, that is expected to be formed when cholesteryl linoleate undergoes oxidation (Estronca, Silva et al. 2012). ChA-treated vascular smooth muscle cells acquire a foam-cell-like phenotype and exhibit membrane fluidity alterations (Alves, Marques et al. 2022). Furthermore, in ChA-treated macrophages the dysfunctional lysosomes full of undigested cargo are exocytic. This outcome can be implicated in the initiation and perpetuation of inflammation in atherosclerosis (Neuza Domingues 2021),(*<u>under revision in the Traffic journal</u>*).

In the present work, we show that ChE are found in the plasma of CVD patients at concentrations considerably higher than in age-matched controls as well as in carotid atheroma plaques. We also demonstrate the role of the most prevalent ChE found in these human tissues, ChA, as a potent inducer of innate inflammatory responses. Importantly, our findings show that ChA changes the inflammatory profile of human monocytes and macrophages. In zebrafish larvae, ChA induces the early features of atheroma formation, such as lipid accumulation and myeloid cell infiltration into their vasculature. Finally, pharmacological approaches indicate that in zebrafish larvae inflammasome as well as the lysosome protease cathepsin B are involved in the inflammatory process. Together, our data indicates that ChA behaves as an endogenous damage associated molecular pattern (DAMP) with inflammatory and pro-atherogenic properties.

## MATERIAL AND METHODS

### Plasma and carotid endarterectomy specimens (CEA)

Blood samples were obtained from CVD patients and healthy individuals after explaining the purpose of the study and obtaining written informed-consent from them or their legal representatives. The entire process was approved by the Ethical Review Board of the Faculty of Medicine of the NOVA University of Lisbon and the Ethics Committee for Health of the Centro Hospitalar de Lisboa Ocidental, Hospital Santa Cruz. All experiments were performed in accordance with the guidelines and regulations. The details for the different cohorts used and sample preparation were already described elsewhere (Gerl, Vaz et al. 2018; Matthiesen, Lauber et al. 2021). After collection, plasma samples were immediately frozen at -80 °C.

Carotid atheroma plaques were isolated from patients with carotid artery disease (CAD) submitted to open endarterectomy at Hospital de Santa Maria / Centro Hospitalar Universitário Lisboa Norte (CHULN). Inclusion criteria were symptomatic carotid artery stenosis of 50-99% or asymptomatic CAD of 70-99% (measured with ultrasound). These samples constitute discarded tissue generated during medical procedures and were collected from patients after informed consent. The excised carotid atherosclerotic plaques were rapidly dissected into maximally diseased atherosclerotic regions and into regions devoid of disease and placed in liquid nitrogen for shotgun lipidomics. Only the necrotic cores of those tissues were processed for shotgun lipidomics. The entire process was approved by the Joint Bioethics Committee of the Faculty of Medicine (University of Lisbon) and Centro Hospitalar Universitário Lisboa Norte (refª 209/18, de 27^th^ of July 2018).

Furthermore, all experiments were performed in accordance with the guidelines and regulations including, the Universal Declaration on Bioethics and Human Rights of UNESCO, 2005; The Charter of Fundamental rights of the EU, 2012; Ethical principles for medical research involving human subjects - Declaration of Helsinki, 2013; EU Regulation 2016/679 and Good Clinical Practice guidelines (Directive 2001/20/EC) and EU Clinical Trials Directive (2005/28/EC). Moreover, they complied with national legislations for the scientific use of human biological samples (Law N° 12/2005 and N° 131/2014).

### Cholesteryl hemiesters synthesis

Cholesteryl hemiesters were prepared following a general procedure described in the literature for the synthesis of ChS (Klein, Kleinman et al. 1974). ChA, was synthesized from the reaction of commercially available cholesterol with freshly prepared azelaic anhydride, which was obtained by reacting azelaic acid with acetyl chloride, as described elsewhere (Hill 1933; Alves, Marques et al. 2022). Cholesteryl hemiglutarate (ChG, cholesteryl-O-(4-carboxybutanoyl)) was prepared by the reaction of commercially available cholesterol with 1.7 molar equivalents of commercially available glutaric anhydride in dry pyridine under reflux for 6 h (Supplementary Figure I). Trituration of the crude product with methanol, followed by purification by flash chromatography with chloroform/methanol/ammonia (50:5:0.25), gave the target ChG as a white solid in 37 % yield. The detailed experimental procedure and characterization data for ChA and ChG are available in the Supplementary Material.

### Lipid extraction, MS lipidomics, data acquisition and analysis

Mass spectrometry-based lipid analysis was performed at Lipotype GmbH (Dresden, Germany) as previously described (Surma, Herzog et al. 2015). Briefly, 50 µL of diluted plasma (equivalent to 1 μL of undiluted plasma) was mixed with 130 μL of 150 mM ammonium bicarbonate solution and 810 μL of methyl tert-butyl ether/methanol (7:2, v/v) was added. 21 µL of an internal standard mixture was pre-mixed with the mixture of organic solvents. The internal standard mixture covered the major lipid classes present in plasma as described previously (Gerl, Vaz et al. 2018). Additionally, 100 pmol per sample of ChS was added for quantification of ChE. The plate was then sealed with a teflon-coated lid, shaken at 4°C for 15 min, and spun down (3000 g, 5 min) to facilitate separation of the liquid phases. One hundred microliters of the organic phase was transferred to an infusion plate and dried in a speed vacuum concentrator. Dried lipids were re-suspended in 40 μL of 7.5 mM ammonium acetate in chloroform/methanol/propanol (1:2:4, v/v/v) and the wells were sealed with an aluminum foil to avoid evaporation and contamination during infusion. All liquid handling steps were performed using a Hamilton STARlet robotic platform with the Anti Droplet-Control feature for pipetting of organic solvents. Samples were analysed by direct infusion in a QExactive mass spectrometer (Thermo Fisher Scientific) equipped with a TriVersa NanoMate ion source (Advion Biosciences). 5 µL were infused with gas pressure and voltage set to 1.25 psi and 0.95 kV, respectively. We scanned for the m/z 400–650 in FTMS − (Rm/z = 200 = 280 000) for 15 s with lock mass activated at a common background (m/z = 529.46262) to detect ChE as deprotonated anions. Automatic gain control was set to 106 and ion trap filling time was set to 50 ms. ChE were identified and quantified by their accurate intact mass from the FTMS spectra with Lipotype Xplorer, a proprietary software developed from LipidXplorer (Herzog, Schwudke et al. 2011; Herzog, Schuhmann et al. 2012). In the CEA samples, for the identification of the various ChE species, 50 mg of tissue was homogenized in 1 mL of MS-water and lipid content of the suspension was extracted as described for the blood plasma.

### Preparation of liposomes

POPC and ChA were mixed at a 35:65 molar ratio. The detailed protocol for liposomes preparation was described previously by our group (Estronca, Silva et al. 2012). POPC liposomes were always used as control.

### Blood-derived monocyte isolation and *in vitro* differentiation into macrophages

Experiments were performed as previously described (Correia, Gaifem et al. 2017), using buffy coats isolated from healthy donors (n = 6) supplied by the Hospital of Braga, after approval of the Competent Ethics Committee (CEC). The human samples received were handled in accordance with the guidelines approved by the CEC. All the donors agreed and signed an authorized consent (ethical approval reference SECVF014/2015). After buffy coats preparation, monocytes were isolated by centrifugation using Histopaque®-1077 (Sigma Aldrich) followed by immunomagnetic separation using a human anti-CD14 purification kit (Miltenyi Biotec). The cell purity was always confirmed by flow cytometry and was superior to 95%. Purified monocytes were then directed used or differentiated *in vitro* into MDM. Cells were cultured in RPMI 1640 medium containing heat-inactivated fetal bovine serum (10%, FBS, Gibco), L-glutamine (2 mM, Gibco), penicillin (50 U mL^−1^, Gibco), streptomycin (50 µg mL−1, Gibco) and HEPES (10 mM, Gibco), and supplemented with the human macrophage colony stimulating factor (20 ng/mL, M-CSF, Peprotech) for 7 days at 37 °C under a humidified 5% CO_2_ air atmosphere. On the third day, new medium was added to the cell culture in order to supply them with new nutrients. In experiments where it was tested the ChA effect on monocyte differentiation into macrophages, the incubation media with M-CSF was supplemented with sub-toxic concentrations of ChA (10 or 25 µM). After differentiation, cell culture was removed for cytokine quantification by ELISA using commercially available kits (Biolegend) and according with manufacturer’s instructions.

### Isolation of neutrophils and culture

Neutrophils were isolated from the peripheral blood collected on EDTA tubes of 5 healthy donors. To separate the neutrophils, whole blood was layered on top of a solution of Histopaque H1117 and H1009 (Sigma Aldrich) and centrifuged for 20 min at 2000 rpm with brake off. After centrifugation, the neutrophils layer was collected to a new tube. Red blood cells (RBC) lysis was performed with RBC lysis buffer (Biolegend) for 15 min at room temperature. Isolated neutrophils were then washed once with Hank’s Balanced Salt Solution (HBSS, Gibco) and cultured on a 96 well plate with U bottom (Sigma Aldrich) in RPMI-1640 (Gibco) supplemented with 1 % autologous plasma. Neutrophils were maintained 4 hours in culture with Brefeldin A (Biolegend) to stop exocytosis.

### Flow Cytometry

MDM were detached by incubation with TrypLE™ Express solution (Life Technologies) at 37 °C for 10 min. For the analysis of surface markers, monocytes and MDM were incubated for 20 min with saturating concentrations of monoclonal antibodies against HLA-DR (Biolegend; clone L243), CD86 (Biolegend; Clone IT2.2), CD206 (Biolegend; clone 15-2) and CD163 (Biolegend; clone GHI/61). Cells were also stained with dihydrorhodamine 123 (DHR 123) and 4-Amino-5-Methylamino-2’,7’ Difluorofluorescein Diacetate (DAF-FM), all from Invitrogen.

For neutrophils experiments, cells were stained with BD HorizonTM Fixable Viability Stain 450 (BD Biosciences), followed by the surface monoclonal mouse anti-human conjugated antibodies: anti-CD15-PE (clone H198) and anti-CD11b-FITC (ICRF44) from Biolegend. Intracellular staining was performed after fixation and permeabilization with Fix/Perm kit (eBiosciences), with the antibodies anti-IL-1β-FITC (clone JK1B-1), anti-IL-6-APC (MQ2-13A5), anti-TNFα-APC (Mab11) from Biolegend and anti-IL-10-FITC (BT-10) from eBioscience.

### Zebrafish maintenance and feeding

The animal study was reviewed and approved by Animal User and Ethical Committees at NMS Research – NOVA Medical School and the Portuguese National Authority for Animal Health (DGAV). Zebrafish lines were maintained in a re-circulating system with a 14 h/day and 10 h/night light cycle at 28°C. Wild-type (AB), *Tg*(*fli1:EGFP)*, *Tg(pu.1:EGFP)* and *Tg(mpeg.mCherryCAAX SH378, mpx:EGFP i114*), the latter was obtained from the Sheffield University. Zebrafish embryos were obtained by *in vitro* fertilization and kept in E3 zebrafish embryo medium at 28°C until reaching the desired developmental stage. Zebrafish larvae were fed twice a day for 10 days, starting at the fifth day post-fertilization (dpf), with control diet (from Mucedola), FC- (normal food supplemented with FC) and ChA-enriched diet (normal food supplemented with 3% ChA w/w, or 52 μmol per gram of food). The food was prepared as previously described (Stoletov, Fang et al. 2009; Domingues, Estronca et al. 2017). For evaluation of vascular lipid accumulation, the food was also supplemented with 10 μg/g red fluorescent CholEsteryl BODIPY® 542/563 C11 (Molecular Probes).

For the experiments with caspase-1 and cathepsin B inhibitors, six dpf SH378 zebrafish larvae were fed overnight with normal diet, FC-enriched (normal food supplemented with 10% FC, w/w, corresponding to 260 µmol per gram of food), ChA-enriched diet (normal food supplemented with 14% ChA w/w, corresponding to 260 µmol per gram of food). Caspase-1 inhibitor 50 µM (c-YVAD-AOM, Calbiochem) and 50 µM cathepsin B inhibitor (ZRLR, Poland kindly given by Ewa Wieczerzak, Department of Biomedicinal Chemistry Faculty of Chemistry, University of Gdansk) were added into the E3 embryonic media 1h before feeding and were present throughout the experiment.

### Lipids microinjection

ChA, FC and POPC liposomes were diluted to the desired concentrations in sterile PBS to be injected in the circulation of zebrafish larvae. POPC liposomes were used as control. The microinjection plate was a petri dish filled with two layers of 2% agarose, one flat to act as a base for the second layer which has gutters on the surface, small trenches to hold the larvae in place. The needles were provided by the fish facility staff. In these, we inserted 2,5 µL of the lipid emulsion. To calibrate the volume of liquid that would be injected, we used a slide marked with a micro-scale and a drop of oil. The tip of the needle was broken with forceps until the drop of the liquid in the oil had the desired diameter. Larvae were then anesthetized with MS-222 (Tricaine) and placed in the petri dish with the gutters. The larvae were aligned in a way that the head was facing away from the user and the dorsal side was facing the needle. For the injection, the needle at around 30° angle was inserted slowly in the posterior cardinal vein, which leads directly to the heart, and pressed the pedal one time to inject the solution (1.5 to 2 nL). The injection was successful when a noticeable flow of fluid increase was observed in the heart.

### Cell Sorting

Larvae were sacrificed on ice to maintain cell viability and transferred to PBS containing 0.1 mg/mL liberase. Tissues were digested at 28 °C for 20-30 min and were mechanically dissociated by resuspending the tissue every 5 min. Cell suspension was passed through a CellTrics 30 µm filter directly to a new Eppendorf and heat inactivated-FBS was added (10 % final concentration). The cell suspension was then sorted in a FACSAriaIIITM high speed cell sorter (Becton Dickinson) equipped with a 561 nm laser (50 mW solid state) and a 630/75 nm BP for mCherry excitation. PBS was used as sheath fluid and run at a constant pressure of 207 kPa with a 100 µm nozzle and a frequency of drop formation of approximately 30 kHz. Cells were directly collected into individual wells of a Nunc™ Lab-Tek™ Chambered plate and allowed to adhere at room temperature for 2 h.

### Lysosomes and neutral lipids staining

After cell sorting, macrophages were incubated for 20 min with BODIPY 493/503 (1:500) and LysoTracker (1:1000) and imaged under a confocal microscope. Alternatively, zebrafish larvae, at 15 dpf, were incubated in a 1:100 dilution of LysoTracker Green in E3 medium at 28.5 °C for 45 min - 1 h, washed twice and then imaged (live imaging).

### Confocal imaging and analysis

At the time points indicated in the figure legends, zebrafish larvae were fixed in 4 % PFA and mounted on a coverslip using mounting media (1 g of DABCO in 10 mL PBS and 40 mL glycerol) prior to imaging. For live imaging, animals were anesthetized with tricaine methanesulfonate and mounted in 1 % low melting point agarose in E3 medium. A Zeiss LSM710 confocal microscope was used and optical sections in the tail area were acquired with a 40x objective.

For quantification of myeloid cells in the caudal vein, fluorescent cells were counted and normalized to the length of the analyzed segment of the vasculature. To quantify lipid accumulation in the vasculature, ImageJ software was used. The lipid structures were delineated and the area was measured. The lipid intensity was normalized to the total area of the vasculature analyzed.

For cell live imaging, cells were imaged in Nunc™ Lab-Tek™ Chambered plate using the water 40x objective. The area and intensity of lysosomes and neutral lipids was measured using ImageJ software.

### Quantitative RT-PCR

Total RNA was extracted with Trizol reagent according to manufacturer instructions. Reverse transcription was performed using the NZY first-strand cDNA synthesis kit (NZYtech). Quantitative PCR was performed in a 96-well optical plate using SYBR green master mix (NZYtech). PCR and data acquisition was performed using the AB7300 Real-Time PCR thermal cycler with *Step One software* (v2.2.2; Applied Biosystems). *eef1al1* - *eukaryotic elongation factor 1 alpha like 1* and *rpl13a* - *ribosomal protein L13a were* used as housekeeping genes to normalize the mRNA expression levels. Target gene expression was determined by relative quantification (ΔΔCt method) to the housekeeping reference genes and the control sample.

The following zebrafish forward and reverse primers were used:

*v-cam1* (TTGCAGTTGTTTCCCACACG; CCTAACGCGGTCCAGACAAA),

*il-1*β (GTAACCTGTACCTGGCCTGC; AACAGCAGCTGGTCGTATCC),

*il-6* (ACGTGAAGACACTCAGAGACG; CGTTAGACATCTTTCCGTGCTG),

*tnf-*α (CAGGGCAATCAACAAGATGG; TGGTCCTGGTCATCTCTCCA),

*il-10* (GCTCTGCTCACGCTTCCTTC; TGGTTCCAAGTCATCGTTGT)

*eef1a1l1* (CCTTCAAGTACGCCTGGGTGTT; CACAGCACAGTCAGCCTGAGAA),

*rpl13a* (TGACAAGAGAAAGCGCATGGTT; GCCTGGTACTTCCAGCCAACTT)

### Zebrafish survival evaluation

Zebrafish survival was first assessed in 5 dpf larvae fed with different diets for 10 days. Larvae were fed twice a day with different doses of FC or ChA. Forty larvae were placed in each test chamber for each food condition. Dead larvae were removed daily. Larval death was expressed as a percentage of the total number of individuals subjected to a given condition and compared to the control population receiving a normal diet.

### Statistical analysis

Results are presented as the mean ± standard error of the mean (SEM). Statistical significance was assessed by one or two-way ANOVA with a Turkey post-test or t-test. A *p* value of 0.05 was considered to be statistically significant.

## RESULTS

### Cholesteryl hemiazelate is the most prevalent cholesteryl hemiester in plasma of CVD patients and in human carotid atheromata

One of the major modifications of LDL trapped in the arterial intima is the oxidation of their PUFA-cholesteryl esters which results in the accumulation of so-called “core aldehydes” (Kamido, Kuksis et al. 1995; Hoppe, Ravandi et al. 1997; Hutchins, Moore et al. 2011) in the fatty streaks and atheromata. We had previously suggested (Estronca, Silva et al. 2012), that the oxygen-rich environment in the arteries promotes the conversion of “core aldehydes” into ChE. Considering the more polar features of ChE, when compared with FC and colesteryl esters, an increased partition of this lipid from the oxidized lipid core of atheromata into the plasma is expected. We, therefore, started by determining the plasma levels of ChE in CVD patients, by applying quantitative mass spectrometry-based shotgun lipidomic analysis.

In order to obtain reliable quantification of ChE, we infused the synthetic standards ChS, ChG (the cholesteryl hemiester of glutaric acid), and ChA. We observed that this class of lipids ionize efficiently in the negative ion mode as deprotonated anions. Moreover, we were able to establish the MSMS fragmentation of the different ChE species and unambiguously detect and quantify them (Supplementary Figure II A-C). We then assessed whether ChE could be detected and quantified in plasma samples. Increasing amounts of a test plasma sample were extracted together with 100 pmol of ChS, an internal standard, which is not normally a component of plasma, and looked in the high resolution (HR)MS and MSMS spectra for the ChE mass spectrometric “fingerprint”. Different ChE, with differing carbon lengths and degrees of unsaturation were observed. Most importantly, their intensities were correlated with the sample volume (Supplementary Figure III). The dynamic range and limits of quantification were established by “spiking” increasing amounts of ChS in 2 µL of the plasma sample (Supplementary Figure IV). The dynamic range was 500 and the inferior limit of quantification was 0.5 µM.

In plasma samples, two ChE were quantitatively analyzable (concentrations >0.5 µM): ChA and a cholesteryl hemiester of 1,11-undecendioic acid (ChU) (Figure 1A). ChA was quantifiable in 88 %, 66 %, and 37 % of acute coronary syndrome (ACS), stable angina pectoris (SAP), and control cohorts, respectively. The cholesteryl hemiester of 1,11-undecendioic acid was only quantifiable in 64 %, 31 %, and 12 % of the ACS, SAP, and control cohorts, respectively. Thus, ChA was the most prevalent of the two ChE quantitatively detectable in plasma. We also found that ChA concentrations were significantly higher in ACS and SAP when compared to age-matched controls. The mean differences from control were 1.04 ± 0.15 µM for ACS and 0.73 ± 0.16 µM for SAP. Of note, the highest ChA detected concentration was 5.95 µM (Figure 1A).

**Figure 1:**
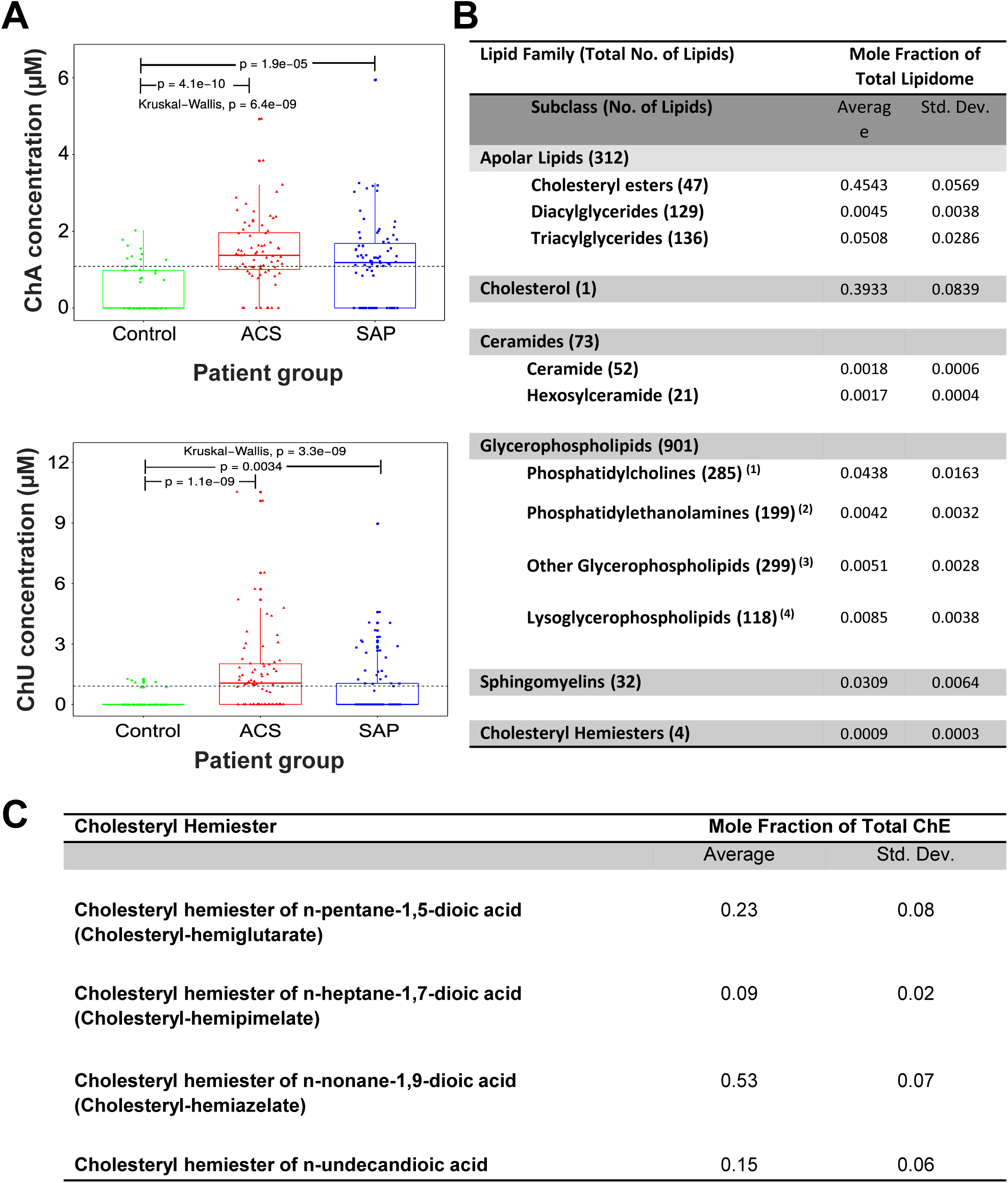
ChA levels are increased in plasma of CVD patients. Boxplots depicting ChA and Cholesteryl hemiester of 1,11-undecendioic acid (ChU) concentrations across patient groups (**A**). Kruskal–Wallis one way analysis of variance was applied to test whether ChA concentration across samples originate from the same distribution. Unpaired two-samples Wilcoxon test was applied as a post hoc test to evaluate significant differences in lipid levels for a patient group versus the control group (indicated by “p”). The horizontal black dashed line indicates the global mean across all samples. Plasmatic concentration of ChA was obtained by shotgun lipidomics of 72 donors with acute coronary syndrome (ACS, including ST-elevation and non-ST-elevation myocardial infarction and unstable angina pectoris), 82 donors with stable angina pectoris (SAP) and 52 age-matched control cohort. **B**. Lipidomes of the atheromata obtained from 6 different carotid artery endarterectomy specimens. ^(1)^ Includes ether Phosphatidylcholines; ^(2)^ Includes ether Phosphatidylethanolamines; ^(3)^ Includes Cardiolipins, Phosphatidic acids, Phosphatidylglycerols, Phosphatidylinositols, and Phosphatidyl serines; ^(4)^ Includes Lysophosphatidic acids, Lysophosphatidylcholines, Ether-Lysophosphatidylcholines; Lysophosphatidylethanolamines, Ether-Lysophosphatidylethanol-amines, Lysophosphatidylglycerols; Lysophosphatidylinositols, and Lysophosphatidylserines. **C**. Cholesteryl Hemiesters detected in atheromata obtained from the six carotid artery endarterectomy specimens.

Shotgun lipidomics was also performed on the lipidic content of human atheromata obtained from human carotid endarterectomy specimens (CEA). By suspending the CEA cores in water, we observed varying degrees of calcification manifested as granular material that could not be easily homogenized. For this reason only the “gruel”-like necrotic core was reliably separated from the rest of the samples. The lipidomic analysis could not, therefore, be related to the mass of atheromata homogenized material. However, our analysis permitted obtaining values of the molar fractions of the total lipid in the “gruel” of the atheromata (Figure 1B). The obtained results show that cholesteryl esters and FC together constitute about 85%, triacyl- and diacylglycerides constitute about 5%, and polar phospholipids constitute about 10% of the total lipid in the “gruel”. We also observed that ChE content of atheromata is almost 1% of the total polar phospholipid content. Additionally, our analysis identified four ChE and their relative proportions are depicted in Figure 1C. ChA constitute a little over 50% of the total ChE content of the atheromata. Interestingly, our findings are well correlated with previous work from other laboratories showing that cholesteryl-9-oxononanoate (the precursor of ChA) was the principal “core aldehyde” in oxidized LDL (Kamido, Kuksis et al. 1993; Kamido, Kuksis et al. 1995) and in atheromata (Hutchins, Moore et al. 2011).

Since our previous results (Estronca, Silva et al. 2012; Domingues, Estronca et al. 2017) have shown that ChE are able to induce “foam cell” formation and inflammation when macrophages and zebrafish larvae are exposed to ChS, we next investigated the pro-atherogenic properties of ChA *in vitro* towards human leucocytes and *in vivo* towards zebrafish larvae.

### The inflammatory profile of human leucocytes is altered after exposure to ChA

In CVD patients, blood stream leucocytes, such as monocytes and neutrophils, are exposed to several bioactive molecules, among them ChA. In this sense, we studied the effect of ChA on monocyte polarization and on their differentiation into macrophages (human monocyte-derived macrophages, MDM). Human monocytes (CD14^+^ cells) were incubated for 24 h with vehicle (POPC liposomes, control) and with sub-toxic concentrations of ChA (ChA:POPC liposomes, 65:35 Molar ratio), followed by analysis of specific cell surface markers by flow cytometry. Surface expression levels of HLA-DR and CD86, markers of classically-activated monocytes, were not affected by the lipid (Figure 2A and B). However, alternatively activated surface markers such as CD206 and CD163 were affected by ChA treatment, with ChA-treated monocytes expressing higher levels of CD206 and lower levels of CD163 when compared with control cells (Figure 2C and D).

**Figure 2.**
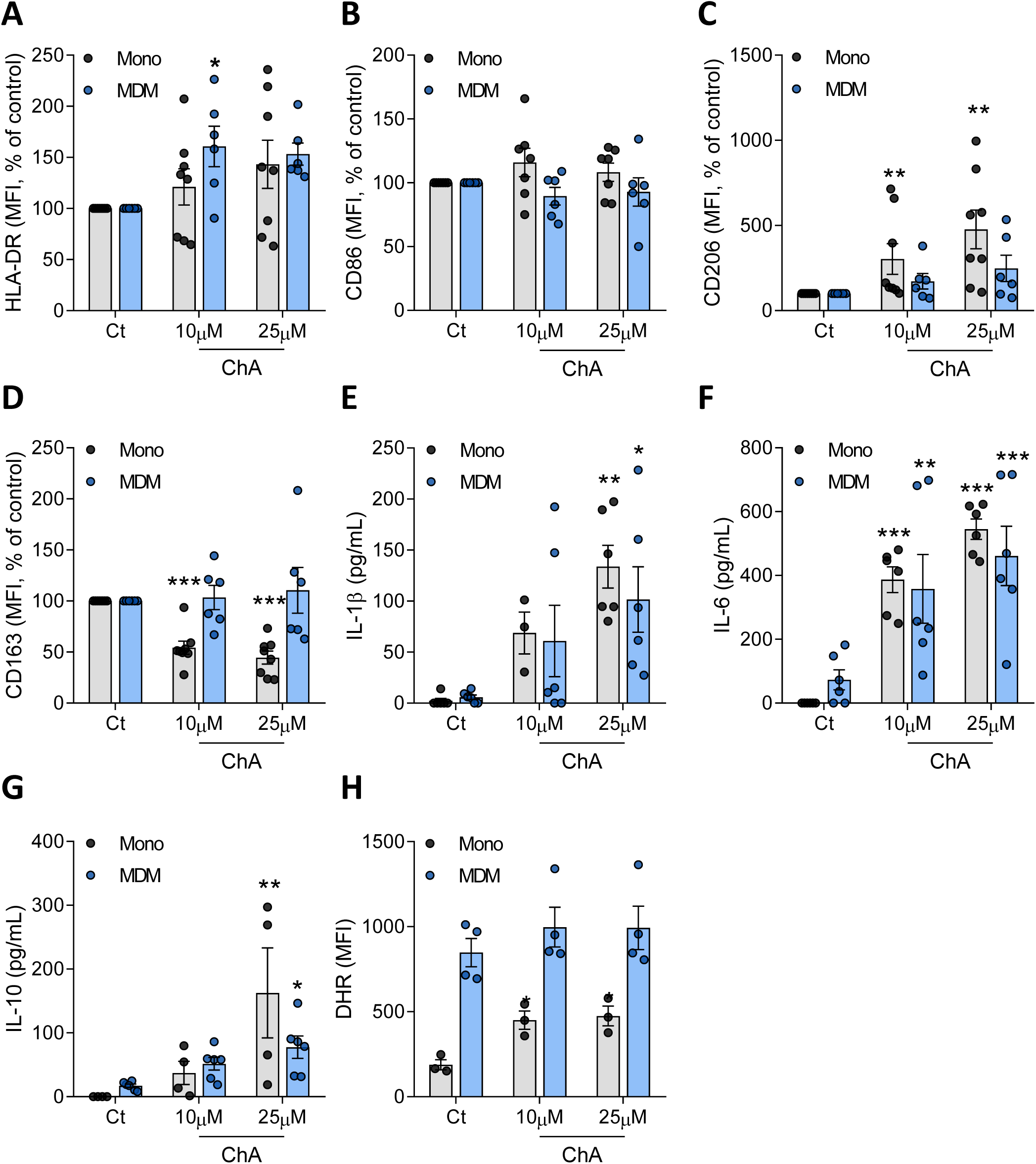
ChA changes the expression of surface markers and the inflammatory profile of human primary monocytes (Mono) and monocyte-derived macrophages (MDM). Peripheral blood was harvested from human volunteers. Mononuclear leukocytes (PBMCs) were isolated from whole blood using histopaque-1077. CD14^+^ cells were purified from total PBMCs by immunomagnetic separation. CD14^+^ monocytes were cultured in the presence of ChA (10 μM or 25 μM) or vehicle (control) for 24 h (n=8) or for 7 days to obtain MDM (n=6). Monocytes were differentiated using rM-CSF. After lipid treatment, a representative sample of cells was stained with monoclonal antibodies to determine surface expression of HLA-DR (**A**), CD86 (**B**), CD206 (**C**) and CD163 (**D**); data represent the mean fluorescence intensity (MFI) ± SEM for all subjects normalized to the control cells. Cell culture supernatants were analysed by ELISA for IL-1β (**E**), IL-6 (**F**) and IL-10 release (**G**). Values represent the mean ± SEM of cytokine released (in pg/mL). Reactive oxygen species production by monocytes and MDM exposed to ChA (**H**). *, *p* < 0.05; **, *p* < 0.01; *** *p* < 0.001.

Next, we studied the impact of ChA on the differentiation of human monocytes into macrophages. For this purpose, human monocytes were incubated with M-CSF to induce their differentiation into macrophages in absence (control) or presence of ChA at sub-toxic concentrations (Supplementary Figure V A). The percentage of CD14^+^CD16^+^ double positive cells after differentiation was evaluated (Supplementary Figure V B). ChA did not affect the monocyte differentiation yield, with the percentage of CD14^int/+^CD16^+^ cells being similar to that found for control cells. Additionally, MDM in the presence of ChA revealed an increased expression of HLA-DR and no differences in CD86 when compared with control cells. These proteins are both surface markers of activated (M1) macrophages. In addition, the surface levels of CD163 CD206, two alternatively activated (M2) markers, revealed no difference in MDM differentiated in presence of ChA (Figure 2C and D).

To further characterize the activation stage of monocytes and MDM exposed to ChA, their cytokine profile was evaluated by ELISA. The results showed in Figure 2E-G demonstrate that ChA-treated monocytes and MDM secreted higher levels of IL-1β, IL-6 and IL-10 than the control cells. Finally, the oxidative burst induced by ChA-loaded in immune cells was quantified through the measurement of DHR fluorescence, a reactive oxygen species (ROS) indicator. The results in Figure 2H demonstrate an increase in ROS production in ChA-treated monocytes, compared with control cells, while in MDM no changes were observed.

Together, these results showed that ChA significantly impacts the monocyte inflammatory phenotype and interferes with the differentiation process of monocytes into macrophages by increasing the expression of inflammatory surface markers, the secretion of inflammatory cytokines and ROS production.

Neutrophils have been implicated in early and advanced atherosclerotic lesions (Doring, Drechsler et al. 2015). Thus, next we assessed the impact of ChA on human neutrophils inflammatory surface marker, CD11b and cytokine production. Human neutrophils were exposed to sub-toxic concentrations of ChA (Supplementary Figure VIA) or vehicle (POPC, control) for 4 h. After incubation, cells expressing the surface marker CD11b and the cytokines IL-1β, IL-6, TNF-α and IL-10, were assessed by flow cytometry. Although, the results presented in Figure 3A showed that the percentage of positive cells for CD11b was not altered by ChA, the percentage of neutrophils positive for IL-1β, IL-6, TNF-α and IL-10 increased in a ChA concentration-dependent manner (Figure 3 B-E), suggesting that ChA activated neutrophils, leading to a more inflammatory profile. This was reinforced, by the observation that IL-1β expression levels, determined when the flow cytometry data was analysed considering the MFI, also increases for the highest concentration of ChA (Supplementary Figure VI C). The observation, that CD11b surface levels in ChA-treated neutrophils decreased, in a dose dependent manner (Supplementary Figure VIB), when the analysis was performed considering the MFI, may be due to the fact that the quantity of this integrin, at the cell surface, is affected by ChA-induced alterations in the fluidity of the membrane lipid bilayer.

**Figure 3.**
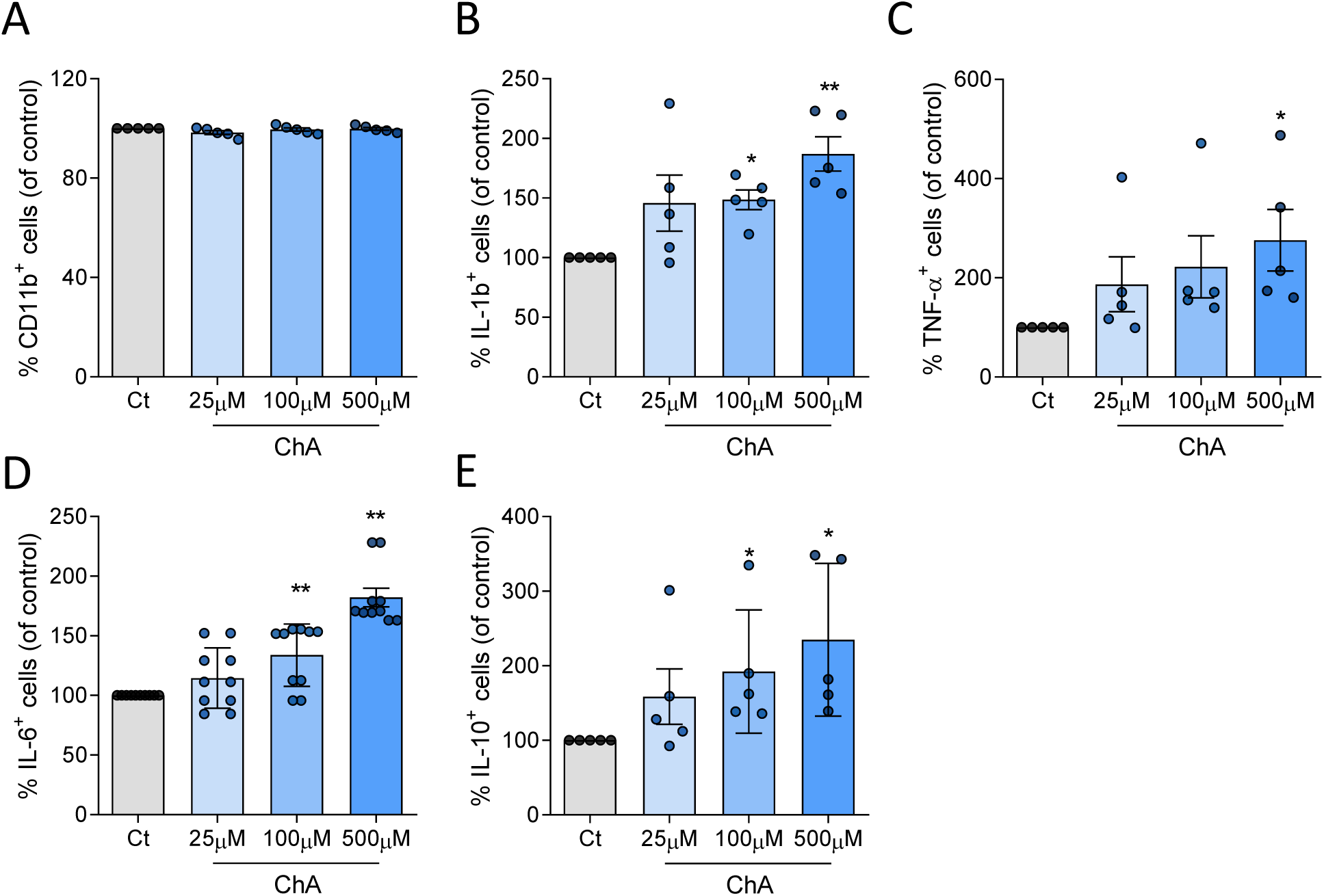
ChA affects the cytokine profile of neutrophils. Neutrophils were harvested from human volunteers (n=5) using a gradient with histopaque-1177 and 1009. Cells were cultured in the presence of ChA (25, 100 and 500 μM) or vehicle (control) for 4 h. After treatment, neutrophils were stained with monoclonal antibodies to determine the expression of CD11b (**A**) and cytokines: IL-1β (**B**), TNF-α (**C**), IL-6 (**D**) and IL-10 (**E**); data represent the percentage of positive cells for each inflammatory marker normalized to the control cells. *, *p* < 0.05; ** *p* < 0.01.

Nevertheless, although ChA seems to activate neutrophils, it is important to have in mind that these *in vitro* assays have to be performed in short time, because neutrophils are short-living cells. Thus, only concentrations 500 times higher than those found in plasma of CVD seem to affect these cells.

### ChA triggers inflammation in zebrafish larvae

Previous work has shown that zebrafish is a suitable animal model to study high-cholesterol diet-induced accumulation of lipids and inflammation (Stoletov, Fang et al. 2009; Fang, Green et al. 2011). Accordingly, the proteins involved in the transport of dietary fat and inflammatory pathways in zebrafish are conserved relatively to mammals (Stein, Caccamo et al. 2007; van der Vaart, Spaink et al. 2012; Progatzky, Sangha et al. 2014). Thus, taking all this information into account we decided to pursued our experiments using this animal model to further characterize the inflammatory effects of ChA.

Since one of the hallmarks of inflammation in atherogenesis is the infiltration of innate immune cells into the arterial intima, we assessed whether ChA, was sufficient to induce infiltration of myeloid cells into the zebrafish larvae vasculature (Figure 4A). Zebrafish larvae feeding started at 5 days post-fertilization (dpf), and as controls we used fish fed with a normal diet (negative control) and a diet enriched in FC (positive control). *PU.1-EGFP* transgenic zebrafish, whose myeloid cells express GFP, fed for 10 days with FC- or ChA-enriched diets, showed, in a dose –dependent manner, an increase in GFP-labelled cells in the caudal vein in comparison with control animals (visualized in Supplementary Figure VIIA and quantified in Figure VIIB). Considering that myeloid cells recruitment with 3% (w/w) of ChA-enriched diet was statistically significant when compared with control we decided to pursue our experiments using this concentration of lipid. Of note, 3% ChA and 2% FC correspond to the same number of moles of these lipids in the supplemented diet (52 µmol/g of food). As supported by our results (Supplementary Figure VII) and as previously described (Stoletov, Fang et al. 2009; Fang, Green et al. 2011; Domingues, Estronca et al. 2017) 4% (w/w) FC (103 μmoles per g of food)-supplemented diet was used as positive control.

**Figure 4.**
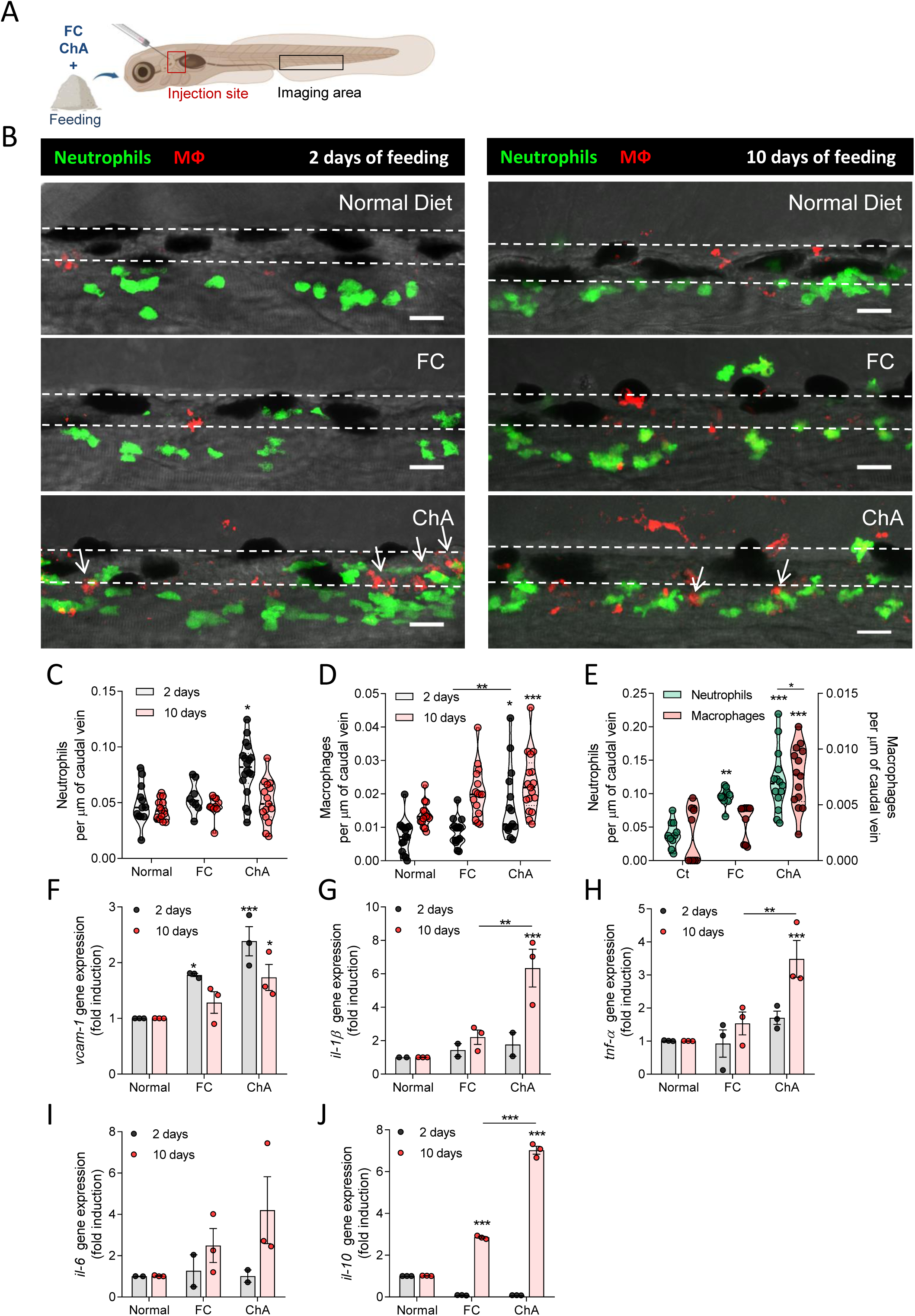
ChA induces neutrophils and macrophages infiltration in zebrafish caudal vein. **A.** Schematic representation indicating with rectangles the sites of lipid microinjection and imaging. **B.** Representative images of the caudal vein of Tg (*mpeg.mCherryCAAX SH378, mpx:EGFP i114*) zebrafish larvae fed for 2 or 10 days with normal, free cholesterol [FC, 4% (w/w), 103 μmoles per g of food, positive control] - or ChA [3% (w/w), 52 μmoles per g of food]- enriched diets and imaged by confocal microscopy. The feeding started at 5 days postfertilization. Images are Z-stacks of fluorescent green (neutrophils) and red (macrophages) cells and the respective bright field. Arrows point to neutrophils and macrophages that are in the close proximity. Dashed lines delineate the caudal vein. Scale bars, 20 μm. **C-D.** Quantification of neutrophils (green cells, **C**) and macrophages (red cells, **D**) after 2 (gray color) or 10 days feeding (red color). Infiltration of neutrophils occurs earlier than macrophages. The results are shown as mean ± SEM of three independent experiments (in each independent experiment at least 5 larvae were analyzed per condition); **E.** Recruitment of neutrophils (green color) and macrophages (red color) into the vasculature of Tg (*mpeg.mCherryCAAX SH378, mpx:EGFP i114*) larvae 24 h after the microinjection of POPC (27 µM, vehicle, Ct), FC:POPC (50 µM FC) and ChA:POPC liposomes (50 µM ChA). The graphs represent the mean ± SEM of three independent experiments (at least 3 larvae per each independent experience were analyzed per condition). **F-J** Quantitative RT-PCR for *vcam-1* (**F**), *il-1*β (**G**), *tnf-*α (**H**), *il-6* (**I**) and *il-10* (**J**) after 2 or 10 days of feeding with normal, FC- or ChA-enriched diets. The values represent the mean ± SEM of three independent experiments (n = 20-30 zebrafish larvae per group); *, *p* < 0.05 **; ***, *p* < 0.001.

To further investigate the kinetics and the type of myeloid cells infiltration, we took advantage of the transgenic zebrafish line mpeg.mCherryCAAX SH378, mpx:EGFP i114, that exhibits neutrophils in green (GFP) and macrophages in red (mCherry). As shown in Figure 4B and quantified in Figure 4C and D, at 2 days of ChA feeding, neutrophils and macrophages infiltration was significantly increased (*p* < 0.05) in ChA-fed animals when compared to FC-fed and normal diet fed animals. At 10 days post-feeding, the number of neutrophils was similar in all experimental conditions (Figure 4C), while the number of macrophages increased in ChA-fed animals in comparison with the 2 days feeding (Figure 4D). These results suggest that the infiltration of neutrophils was transient while the infiltration of macrophages persisted through time.

Since the concentration of lipid in circulation is not controlled during the animal feeding process, we decided to confirm the results obtained above by injecting directly the lipid into the fish blood stream. At 24 h after ChA and FC microinjection, the number of neutrophils in zebrafish caudal vein increased significantly. At this time point, an increase in number of macrophages was also observed for ChA-microinjected animals while for FC-treated animal this outcome was not obtained (Figure 4E). These results indicate that neutrophils infiltration occurred earlier than macrophages and that ChA was more inflammatory that FC. Of note, microinjection of embryonic medium (control, Ct) and POPC liposomes (vehicle) did not impact the number of inflammatory cells into the vasculature, suggesting that the increase in the inflammatory cells infiltration was indeed due to FC and ChA (Supplementary Figure VII C - E).

The expression of adhesion molecules can result in the recruitment of inflammatory cells into the vasculature. Therefore, we decided to measure the mRNA levels of the vascular cell adhesion molecule (*vcam-1*) in zebrafish larvae after feeding. The results revealed an increase in *vcam-1* gene expression at 2 and 10 days of ChA feeding (Figure 4F), which was in accordance with the accumulation of myeloid cells in larvae vascular walls. To gain more insights about the inflammatory effects of ChA using an *in vivo* system, we next investigated the impact of ChA in the mRNA levels of inflammatory cytokines. Initially, we confirmed the ability of zebrafish larvae to express cytokines at 15 days post-fertilization, by incubating zebrafish with LPS, as previously reported (Progatzky, Sangha et al. 2014). An overnight treatment of zebrafish larvae with LPS induced a significantly increase in *il-1*β, *tnf-*α and *il-6* (as well as in *vcam-1*) mRNA expression as compared to untreated control zebrafish (Supplementary Figure VII F-I). Next, we analyzed the levels of these cytokines in larvae fed for 2 and 10 days with normal, FC- and ChA-enriched diets. At 2 days post-feeding, no statistically significant differences were observed in gene expression levels for the inflammatory cytokines tested (Figure 4 G-J). However, at 10 days post-feeding, the results obtained showed an increase on *il-1*β (Figure 4G), *tnf*-α (Figure 4H) and *il*-6 (Figure 4I) expression in ChA-fed larvae. We also observed alterations in *Il-10* (Figure 4J) mRNA levels induced by FC- and ChA-enriched feeding. Thus, these results obtained using a simple *in vivo* model pointed to an inflammatory behavior of ChA.

Recently, we showed that ChA-treated murine macrophages exhibit features of foam cells, namely enlarged dysfunctional lysosomes and exuberant neutral lipid accumulation (Neuza Domingues 2021) (*<u>under revision in the Traffic journa</u>l*). Thus, we questioned the capacity of ChA to induce neutral lipid accumulation in zebrafish macrophages subjected to ChA feeding for 10 days. The mCherry positive macrophages were isolated by fluorescence-assisted cell sorting (FACS) and stained for neutral lipids with BODIPY 493/503. As shown in Figure 5A and quantified in B, neutral lipid accumulation in macrophages isolated from larvae fed with ChA was much higher than in larvae fed with FC or with a normal diet (macrophages are outlined by dashed lines). Therefore, we next considered whether these macrophages from lipid enriched diet fed animals would also show evidence of lysosomal dysfunction. Since changes in lysosome area are an indication of their malfunction, we decided to quantify this parameter in macrophages isolated from larvae fed with the different diets by using Lysotracker, a lysosomotropic dye. Remarkably, the results shown in Figure 5C and quantified in Figure 5 D demonstrated an increase in lysosomal area in macrophages isolated from zebrafish larvae fed with a ChA-enriched diet, compared with the controls. To confirm this latter result, we also evaluated the lysosomal morphology in macrophages in live larvae. As shown in Figure 5E macrophages in the vasculature of zebrafish fed with a ChA-enriched diet presented enlarged lysosomes. Interestingly, lysosomal positioning within macrophages from larvae subject to the different types of diet was also impacted. In macrophages of FC-fed animals, lysosomes seemed to be clustered in the perinuclear region while lysosomes in macrophages from ChA-fed animals were located more proximal to the macrophage edges.

**Figure 5.**
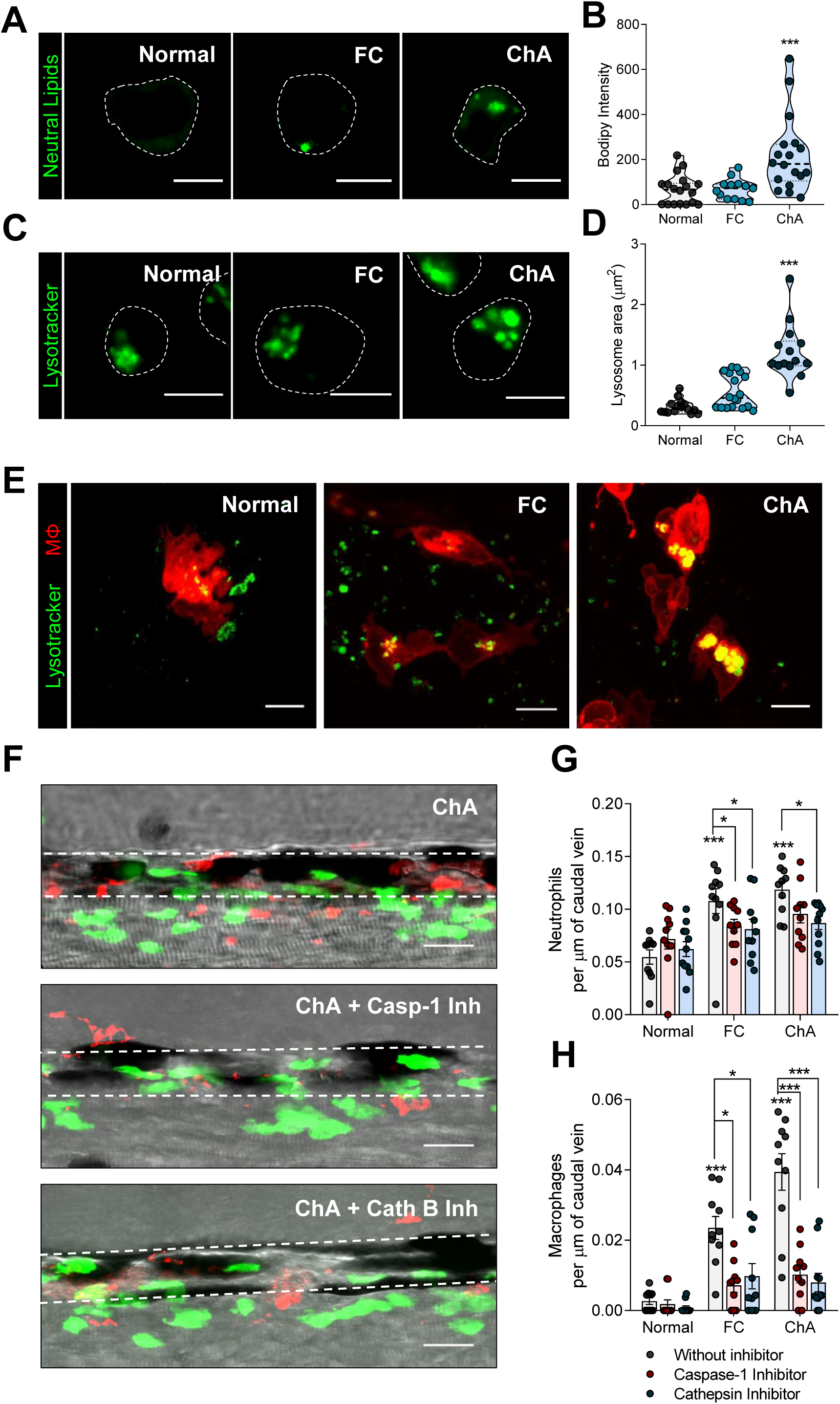
Inflammasome and cathepsin-B inhibition decrease ChA-induced myeloid cell infiltration. **A-D.** Zebrafish larvae were fed for 10 days with normal, 4% FC- or 3% ChA-enriched diets and macrophages were isolated by Fluorescence Activated Cell Sorting (FACS). **A.** Representative confocal images of macrophages stained with BODIPY 493/503 to visualize neutral lipids. **B.** Quantification of neutral lipid accumulation on isolated macrophages**. C.** Representative confocal images of macrophages stained with lysotracker to visualize lysosomes/acidic organelles. Scale bars, 5 μm. **D.** Quantification of the lysosome area on isolated macrophages. In **B** and **D**, the results are the mean ± SEM of two independent experiments (at least 10 cells were analyzed per condition); ***, *p* < 0.01. **E.** *In vivo* lysosomal imaging of macrophages (m-Cherry) from larvae fed with different diets. Lysosomes were stained with lysotracker (green). Scale bars, 5 μm. **F-H**. Effect of Caspase 1 (Casp-1) and Cathepsin B (Cath B) inhibitors on neutrophils and macrophages infiltration into the caudal vein of Tg (*mpeg.mCherryCAAX SH378, mpx:EGFP i114*) zebrafish larvae fed with normal, 10% FC- or 14% ChA-enriched food. **F.** Representative confocal images. Images are Z-stacks of fluorescent green (neutrophils) and red (macrophages) cells and the respective bright fields. Dashed lines delineate the caudal vein. Scale bars, 20 μm. **G and H.** Quantification of Casp-1 and Cath B inhibitors on neutrophils (**G**) and macrophages (**H**) infiltration into the caudal vein. Three independent experiments were performed and at least 3 larvae were analyzed per experiment (total of >10 larvae per condition).

Several studies have been reported that oxidized lipids from oxLDL, saturated fatty acids and cholesterol crystals activate the NOD-like receptor protein 3 (NLRP3) inflammasome in macrophages, resulting in IL-1β maturation and secretion (Duewell, Kono et al. 2010; Wen, Gris et al. 2011; Sheedy, Grebe et al. 2013; Emanuel, Sergin et al. 2014). Two distinct signals are required for NLRP3 activation: a priming signal, leading to the activation of NF-κB and subsequent upregulation of NLRP3 and pro-interleukin-1β; and an activation signal, that can be provided by a plethora of stimuli that will trigger a series of molecular events culminating in NLRP3 activation. Inflammasome signaling ultimately leads to the activation of the effector caspase-1 that is responsible for the cleavage of pro-IL-1β into IL-1β (Supplementary Figure VIII). The signal 2 can be provided by the presence of cathepsin B, a lysosomal protease, in the cytosol, that occurs when lysosomal membrane integrity is lost. Considering the observed increased on *il-1*β expression (Figure 4G) and the changes on lysosome morphology, we decided to investigate the involvement of the inflammasome as well as the role of lysosomes in our experimental settings. For this purpose, we used caspase-1 and cathepsin B inhibitors. Since lipid-enriched diets led to a significant increase in myeloid cells in the caudal vein of zebrafish, we took advantage of this robust read out to evaluate the role of inflammasome and lysosomes on the inflammation induced by ChA. Zebrafish larvae were pre-treated with caspase-1 and cathepsin B inhibitors for 1 h and then, the animals were fed overnight with high doses of FC- and ChA-enriched diets, 10% (w/w) and 14% (w/w), respectively. Our data showed that a single dose of high amounts of FC and ChA were sufficient to induce a significant increase in neutrophils and macrophages recruitment into the vasculature of zebrafish larvae (Figure 5F-H). Interestingly, treatment either with caspase-1 or cathepsin B inhibitors deeply decreased the recruitment of neutrophils (Figure 5G) and macrophages (Figure 5H) in FC- and ChA-enriched diets in comparison with control conditions (Figure 5 F-H). We also observed that the effect of cathepsin B inhibitor was more pronounced on the infiltration of macrophages in ChA-than in FC-fed zebrafish, which can be correlated with the observed increased in size and intracellular positioning of lysosomes within macrophages.

These results reveal that, as reported for cholesterol (Progatzky, Sangha et al. 2014), ChA induces the activation of caspase-1 and lysosome membrane permeability loss, that together are involved in the activation of inflammatory pathway resulting in myeloid cell infiltration.

### ChA potentiates lipid accumulation at the sites of the vasculature bifurcation and impacts zebrafish survival

Considering the accumulation of neutral lipids in macrophages in ChA-fed zebrafish, we questioned whether neutral lipid accumulation was also observed at the sites of the vasculature bifurcation. We started by analyzing the presence of lipid deposits on the vasculature of AB larvae fed with normal, FC- and ChA-diets supplemented with a red fluorescent cholesteryl ester analog (BODIPY 542/563 C11). The larvae caudal veins were imaged 10 days after feeding. As shown in Figure 6A, ChA- and FC-fed larvae showed many focal areas of bright red fluorescence, which we interpreted as lipid accumulation in the vessel wall of the caudal vein. Larvae fed with a normal diet did not exhibit fluorescent lipid structures. The percentage of lipid structure area of the bright red fluorescent structures in the vessel walls, obtained by confocal Z-stacks of the AB zebrafish larvae, was quantified (Figure 6B). Our data show that both, FC- and ChA-enriched diets increased lipid accumulation in a dose-dependent manner. Remarkably, ChA induced lipid-accumulation values much higher than an equivalent amount of FC, suggesting a more pro-atherogenic activity. Of note, the lipid accumulation was already significantly increased in larvae fed for 5 days with ChA when compared to the control groups (Supplementary Figure IX).

**Figure 6.**
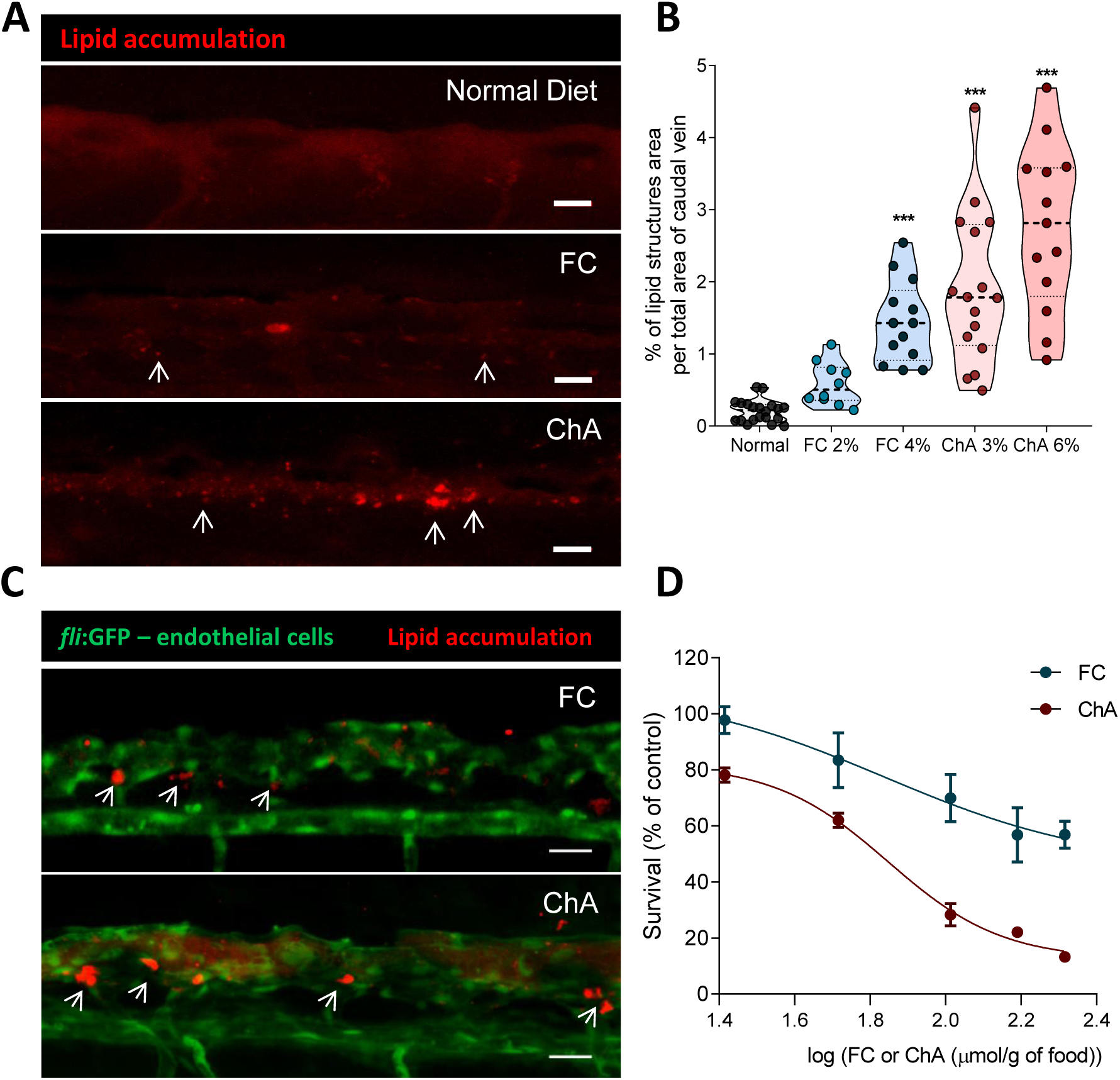
ChA induces lipid accumulation in the vasculature and is toxic to zebrafish larvae. 5 days postfertilization, zebrafish larvae were fed for 10 days with normal (in gray), FC- (in blue), or ChA-enriched diets (in red). **A.** Confocal z-projection images of fluorescent lipid deposits (in red, indicated by the arrows) of caudal vein of AB larvae. For lipid structures visualization, diets (normal, 2 or 4% FC-, 3 or 6% ChA-enriched food) were supplemented with 10 μg/g of a red fluorescent cholesteryl ester. Scale bars, 20 μm. **B.** Quantification of total lipid structures area in the zebrafish caudal vein. Fluorescent images of at least 10 larvae were quantified per condition. The results are shown as mean ± SEM; ***, *p* < 0.001. **C.** Z-projection of the caudal vein of *fli1:EGFP* larvae fed as above. Green fluorescence corresponds to endothelial cells (ECs) and red fluorescence corresponds to deposits of lipids localized at the bifurcation sites (arrows). Scale bars, 20 μm. **D.** Evaluation of larvae survival as a function of FC (in blue) or ChA (in red) concentration in the diet. Lipid concentration is given by logarithm of 26 µmol/g of food (FC 1 %, ChA 1.5 %), 52 µmol/g of food (FC 2 %, ChA 3%), 103 µmol/g of food (FC 4 %, ChA 6%), 155 µmol/g of food (FC 6 %, ChA 9 %) and 207 µmol/g of food (FC 8 %, ChA 12 %). Results are mean of two independent experiments (each experiment with 40 larvae). The error bars indicate the SEM.

To further characterize lipid accumulation in the vasculature, *fli1:EGFP* larvae which constitutively express GFP in the endothelial cells, were fed with different lipid enriched-diets and the lipid tracking probe as described above. We observed that the fluorescence lipid accumulation was subendothelial and was localized at sites of blood vessel bifurcation (Figure 6C). This subendothelial localization is well correlated with what has been described in humans and mice (Ku, Giddens et al. 1985; Tabas, Garcia-Cardena et al. 2015).

Finally, we addressed the impact of ChA-enriched diet on zebrafish larvae survival. For this purpose, zebrafish were exposed to different doses of ChA and FC- for 10 days. After this period of time, viable larvae were counted, and the survival was normalized to the data obtained with animals fed with a normal diet. The results depicted in Figure 6D showed that ChA was toxic to zebrafish in a dose-dependent manner. For all doses tested, we also observed higher toxicity of ChA compared with FC. Using the Hill equation, we determined an ED50 of 61 and 121 µmol/g, with the minimum percentage of survival (Ymin) of 12% and 46%, for ChA and FC, respectively.

Together our results demonstrated that ChA was able to induce lipid accumulation in zebrafish vasculature decreasing the survival of the animals.

## DISCUSSION

Several ω-oxoester precursors of ChE (mono-esters of C4 through C9 diacids) of cholesterol have been identified in oxLDL (Kamido, Kuksis et al. 1993) and in the “core aldehyde” fraction of *ex vivo* samples of human atheromata (Hutchins, Moore et al. 2011). These oxoesters are expected to be further oxidized in extracellular as well as intracellular environments to their corresponding ChE (Estronca, Silva et al. 2012). When formed, ChE may be expected to accumulate at the polar surface of the LDL particles, conferring a negative surface charge. They would also partition via passive diffusion and trans-membrane translocation into all membranes and the cytosol of neighboring cells. Considering that cholesteryl linoleate, the predicted precursor of ChA, is the most abundant cholesteryl ester in plasma (Matthiesen, Lauber et al. 2021), we estimated that ChA would be one of the main end-products of cholesteryl linoleate oxidation and increased in the plasma of CVD patients. It is, in fact, the most prevalent ChE in plasma of CVD patients and atheromatous plaque material obtained from human endarterectomy specimens. The plasma levels of ChA are, on the average, about 3-fold higher in CVD patients than in controls. As far as we know, this is the first time that these lipids have been detected and reported to be increased in the plasma lipidome of CVD patients. Here, we show that this newly identified lipid affects the (i) inflammatory profile of human monocytes, human monocytes-derived macrophages and neutrophils; (ii) promotes recruitment of macrophages and neutrophils into the vasculature of zebrafish larvae; (iii) induces lipid accumulation in the vascular wall, and (iv) results in a decrease of the animal’s lifespan.

Monocytes, monocytes-derived macrophages and neutrophils treated with ChA exhibit different surface markers profiles when compared with control cells. In our experimental settings neither monocytes nor macrophages acquire the classically Th-1 polarized M1 or alternatively Th-2 polarized M2 monocytes/macrophage phenotypes (Auffray, Sieweke et al. 2009; De Paoli, Staels et al. 2014; Gibson, Domingues et al. 2018). These findings suggest a distinct monocyte and macrophage phenotype from the ones already described in literature. A similar statement can be made for neutrophils. Furthermore, beyond the fact that ChA-driven inflammatory profile in neutrophils is dose-dependent, the concentrations of ChA necessary to observe an inflammatory response were much higher than those required for monocytes or macrophages. This can be explained on the one hand, by the fact that neutrophils are short lived. On the other hand, neutrophils could be more resistant to ChA. Indeed, activation of neutrophils, the main type of leucocytes in the blood stream, by low ChA concentrations in the blood stream could be devastating to the CVD patients due the neutrophil extracellular trap (NET) formation (netosis) and subsequently thrombus formation (Liu, Carmona-Rivera et al. 2018; Dou, Kotini et al. 2021).

Common to the human leukocytes studied and to the zebrafish larvae is the fact that ChA increases the proinflammatory cytokine IL-1β, a crucial cytokine implicated in the initiation and development of atherosclerosis (Chi, Messas et al. 2004; Baldrighi, Mallat et al. 2017). This is in agreement with the significant 15% reduction in risk of myocardial infarction and stroke seen in the CANTOS clinical trial that targeted IL-1β with neutralizing antibodies in CVD patients (Ridker, Everett et al. 2017). Thus, ChA could be considered to be a newly identified DAMP contributing to the inflammatory activity of oxLDL (Choi, Sviridov et al. 2017; Miller and Shyy 2017). This finding is important since in contrast to oxPL, oxidized cholesterol, and some other oxidized cholesteryl ester (oxCE) products (Choi, Harkewicz et al. 2009; Oskolkova, Afonyushkin et al. 2010; Miller, Choi et al. 2011; Choi, Yin et al. 2013; Ravandi, Leibundgut et al. 2014; Choi, Sviridov et al. 2017; Miller and Shyy 2017; Gonen and Miller 2020), ChE have been almost completely ignored in the literature.

*In vivo* ChA triggers neutrophils and macrophages infiltration into the vasculature of zebrafish larvae (Figure 7). This is a key event in the pathogenesis of atherosclerosis since their recruitment may result in the exacerbated oxidation of LDL, setting the stage for catalytic expansion of the atherosclerotic lesion and the full-blown spectrum of atherosclerosis (Cynshi and Stocker 2005). In our experimental settings, neutrophils infiltration is transient. On the contrary, macrophages accrue with time within the vasculature. This feature is compatible with a non-resolved inflammation state. Macrophage-accumulation coincides with an increase on IL-1β transcripts and pharmacological inhibition of the inflammasome dampens the inflammatory cells infiltration. This can be explained by the fact that rise in inflammatory mediators leads to additional rounds of monocyte and neutrophils recruitment (Weber and Noels 2011). Additionally, similarly with what has been described in the literature (Duewell, Kono et al. 2010; Emanuel, Sergin et al. 2014; Progatzky, Sangha et al. 2014) lysosome membrane permeabilization seems also to be the culprit of inflammation. Most interestingly, inhibition of the lysosomal cathepsin B activity, leads to a reduction of neutrophils and macrophages in the vasculature, suggesting the involvement of lysosomes in ChA-mediating inflammation by IL-1β production. Thus, we can envision that lipid accumulation in dysfunctional lysosomes causes loss of lysosome membrane permeability, with cathepsin B leakage into the cytosol and subsequent inflammasome activation and IL-1β secretion (Figure 7).

**Figure 7.**
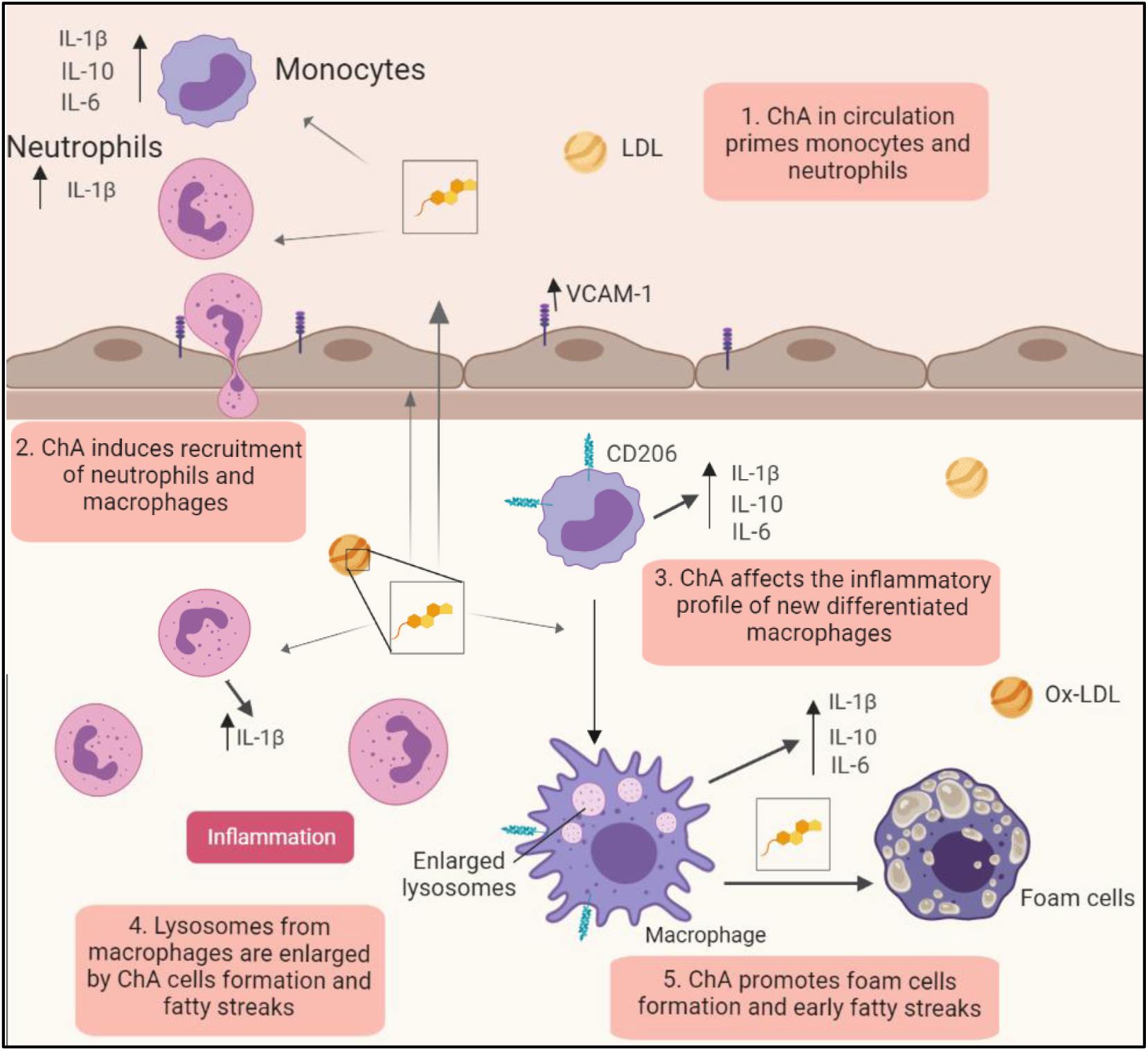
Working model of the pro-atherogenic properties of ChA. ChA is an end product of cholesteryl linoleate oxidation, generated in the arterial intima. Due to its amphiphilic properties, ChA can be detected in the plasma of CVD patients. The presence of ChA in circulation can imprint an inflammatory phenotype in the circulating monocytes and neutrophils (1), conditioning the immunological response in the arterial intima. ChA promotes the recruitment of innate immune cells, neutrophils and monocytes into the vasculature (2). Here, neutrophils in presence of ChA secret IL-1β, which can interfere with monocyte/macrophage priming. Monocytes differentiated in presence of ChA are activated, increasing the secretion of inflammatory cytokines: IL-1β, IL-6 and IL-10. In activated macrophages, ChA induces lipid accumulation (foam cells) and lysosomal dysfunction, conferring then the second signal necessary for IL-1β secretion mediated by inflammasome activation (4). IL-1β can initiate a propagation loop of the inflammation, increasing the macrophages secretion of IL-6 and TNF-α. On another hand, dysfunctional lysosomes will decrease the clearance capacity of macrophages, leading to lipid accumulation in the arterial intima (5).

Vascular inflammation is accompanied by vascular lipid accumulation. Using transgenic zebrafish with GFP-labelled endothelial cells fed with ChA, we observed an increase in lipid accumulation at the caudal vein bifurcations, where turbulent blood flow is expected to occur. Our results are in accordance with those observed for zebrafish fed with FC- and ChS-enriched diets (Stoletov, Fang et al. 2009; Domingues, Estronca et al. 2017), as well as with published data in humans and mice, where a higher propensity of lipid-driven disease is observed in the arteries branches with disturbed flow (Ku, Giddens et al. 1985; Tabas, Garcia-Cardena et al. 2015). Finally, inflammation and lipid accumulation impact zebrafish larvae life expectancy, ChA being even more lethal than FC (Stoletov, Fang et al. 2009).

Thus, in this comprehensive study, we showed *in vivo* and *in vitro* inflammatory and atherogenic properties of ChA, a newly identified and quantified oxidized lipid in CVDs patients. In the future, a better understanding of the molecular mechanisms underlying the biological effects of ChA may identify specific targets for future therapies, offering better focused and potentially personalized treatment of cardiovascular disease. Moreover, a further analysis on the diagnostic and/or prognostic potential of ChA concentration in plasma of patients may provide novel clinical tools to tackle CVD.

## Acknowledgements

We acknowledge the technical support of the Microscopy and Fish Facilities NOVA Medical School. The authors also acknowledge the UC-NMR facility for obtaining the NMR data (http://www.nmrccc.uc.pt).

## Author Contributions

ND: conception and design, acquisition of data, analysis and interpretation of data, article draft

JG: acquisition of data, analysis and interpretation of data

RM: analysis and interpretation of data

DS: acquisition of data

LB: acquisition of data

ARAM: acquisition of data

MI.L.S: acquisition of data

JS: acquisition of data, analysis and interpretation of data

CK: acquisition of data, analysis and interpretation of data

MA.S: acquisition of data, analysis and interpretation of data

MS.A: acquisition of data, analysis and interpretation of data

GR: acquisition of data, analysis and interpretation of data

PA: acquisition of data, analysis and interpretation of data

JF: acquisition of data, analysis and interpretation of data

RGM: acquisition of data, analysis and interpretation of data

LMP: acquisition of data, analysis and interpretation of data

KS: interpretation of data

TMVDPM: interpretation of data

GC: acquisition of data, analysis and interpretation of data

AJ: interpretation of data

RS: acquisition of data, analysis and interpretation of data, revisions in intellectual content of Ms

WV: analysis and interpretation of data, article draft

OVV: conception and design, analysis and interpretation of data, article draft, funding, supervision

