## Supplemental Methods for "Cholesteryl hemiazelate Identified in Cardiovascular Disease Patients Causes *in vitro* and *in vivo* Inflammation"

**SUPPLEMENTARY METHODS**

**Cholesteryl hemiesters synthesis**

**General Remarks:** Thin-layer chromatography (TLC) analyses were performed using precoated silica gel plates. Flash column chromatography was performed with silica gel 60 as the stationary phase. ^1^H NMR spectra were recorded on an instrument operating at 400 MHz and ^13^C NMR spectra were recorded on an instrument operating at 100 MHz. The spectra were recorded in CDCl_3_ as solvent, chemical shifts are expressed in parts per million (ppm) relatively to internal tetramethylsilane (TMS) and coupling constants (*J*) are in hertz (Hz). Infrared spectra (IR) were recorded in a Fourier Transform spectrometer coupled with a diamond Attenuated Total Reflectance (ATR) sampling accessory. High-resolution mass spectra (HRMS) were obtained on an electrospray (ESI) APEX-Qe or microTOF mass spectrometer. Melting points were determined in open glass capillaries. Azelaic acid, acetyl chloride, glutaric anhydride, cholesterol and ammonia were purchased from commercial sources and used as received. Pyridine was dried over KOH. Ethanol and methanol were purified by distillation. Chloroform was distilled and passed through a column of basic alumina.

Cholesteryl hemiesters were prepared following the synthetic pathway outlined in Supplementary Figure I.

**Cholesteryl hemiazelate (ChA, cholesteryl *O*-(8-carboxyoctanoyl)):** Azelaic anhydride was prepared by the reaction of azelaic acid with acetyl chloride. A solution of cholesterol (2.62 g, 6.78 mmol) and azelaic anhydride (3.00 g, 17.63 mmol) in dry pyridine (27 mL) was heated under reflux for 7 h. Pyridine was removed under reduced pressure and the crude product was purified by flash chromatography [chloroform/methanol/ammonia (50:5:0.25, v/v)] giving cholesteryl hemiazelate as a fluffy solid. Further recrystallization from ethanol gave cholesteryl hemiazelate as a white solid (2.16 g, 57%). m.p. 79.5-80.4 ºC. IR (ATR) *ν* 1168, 1234, 1466, 1702, 1730, 2850, 2866 and 2932 cm^-1^. RMN ^1^H (CDCl_3_, 400 MHz) *δ* = 0.68 (s, 3H), 0.86 (d, *J* = 1.7 Hz, 3H), 0.87 (d, *J* = 1.7 Hz, 3H), 0.91 (d, *J* = 6.5 Hz, 3H), 0.94-1.65 (m, 35H), 1.81-1.87 (m, 3H), 1.93-2.02 (m, 2H), 2.25-2.36 (m, 6H), 4.58-4.65 (m, 1H), 5.37-5.38 (m, 1H). RMN ^13^C (CDCl_3_, 100 MHz) *δ* = 11.9, 18.7, 19.3, 21.0, 22.6, 22.8, 23.8, 24.3, 24.6, 24.9, 27.8, 28.0, 28.2, 28.9, 31.9, 31.9, 33.8, 34.6, 35.8, 36.2, 36.6, 37.0, 38.1, 39.5, 39.7, 42.3, 50.0, 56.1, 56.7, 73.7, 122.6, 139.7, 173.3, 179.1. HRMS (ESI) *m/z*: Calcd for C_36_H_61_O_4_ [M+H]^+^ 557.45644; found 557.45615.

**Cholesteryl hemiglutarate (ChG, cholesteryl *O*-(** **4-carboxybutanoyl)):** A solution of cholesterol (3.71 g, 9.60 mmol) and glutaric anhydride (1.90 g, 16.32 mmol) in dry pyridine (35 mL) was heated under reflux for 6 h. Pyridine was removed under reduced pressure and the crude solid product was triturated with methanol. The filtered solid was further purified by flash chromatography [chloroform/methanol/ammonia (50:5:0.25, v/v)] giving cholesteryl hemiglutarate as a white solid (1.79 g, 37%). m.p. 124.4-125.9 ºC. IR (ATR) *ν* 1174, 1282, 1376, 1428, 1702, 1729, 2866 and 2932 cm^-1^. RMN ^1^H (CDCl_3_, 400 MHz) *δ* = 0.68 (s, 3H), 0.86 (d, *J* = 1.6 Hz, 3H), 0.87 (d, *J* = 1.6 Hz, 3H), 0.91 (d, *J* = 6.5 Hz, 3H), 0.94-1.57 (m, 25H), 1.80-1.88 (m, 3H), 1.92-2.05 (m, 4H), 2.30-2.45 (m, 6H), 4.58-4.66 (m, 1H), 5.37-5.38 (m, 1H). RMN ^13^C (CDCl_3_, 100 MHz) *δ* = 11.9, 18.7, 19.3, 19.9, 21.0, 22.6, 22.8, 23.8, 24.3, 27.8, 28.0, 28.2, 31.9, 31.9, 32.8, 33.5, 35.8, 36.2, 36.6, 37.0, 38.1, 39.5, 39.7, 42.3, 50.0, 56.1, 56.7, 74.1, 122.7, 139.6, 172.3, 177.8. HRMS (ESI) *m/z*: Calcd for C_32_H_53_O_4_ [M+H]^+^ 501.39384; found 501.39160.
