## Supplementary material for "Cholesteryl hemiazelate Identified in Cardiovascular Disease Patients Causes *in vitro* and *in vivo* Inflammation": Supplemntal Figures

### SUPPLEMENTARY FIGURES

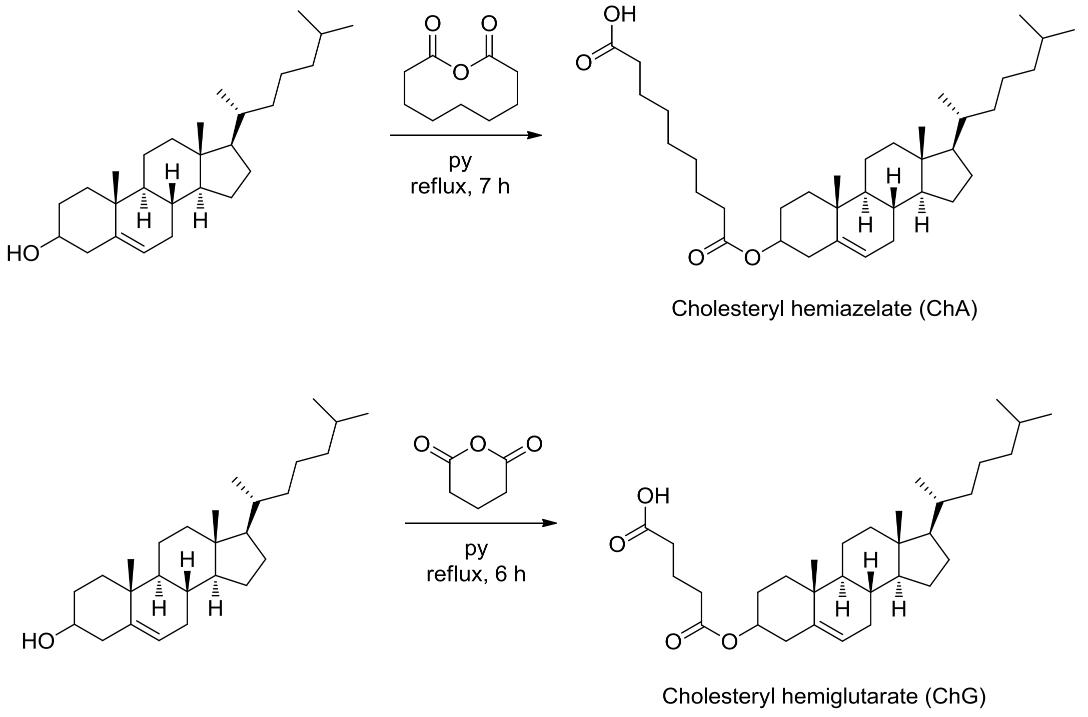


**Supplementary Figure I.** Scheme of the Synthetic strategy towards cholesteryl hemiazelate (ChA, cholesteryl O-(8-carboxyoctanoyl) and cholesteryl hemiglutarate (ChG, cholesteryl O-( 4-carboxybutanoyl) ).

### Supplementary Figure II

A

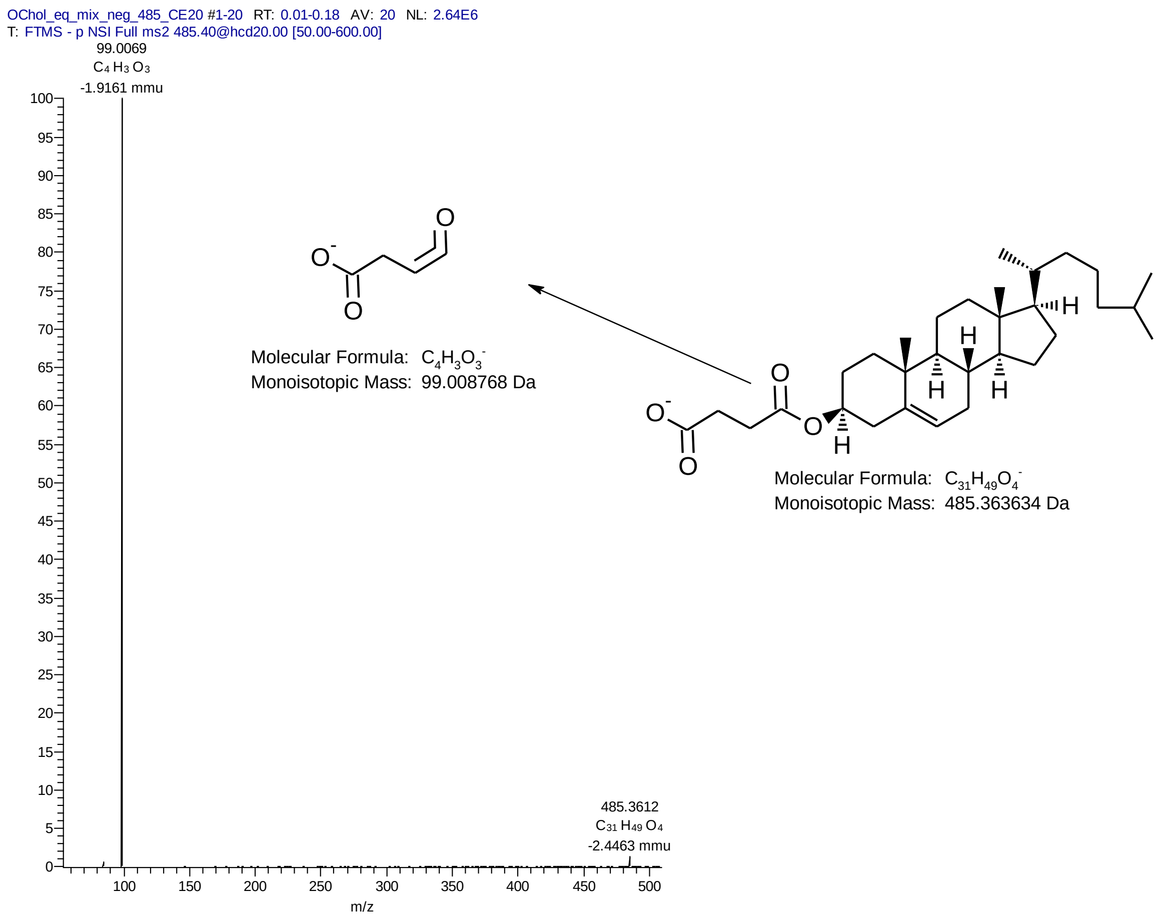


B

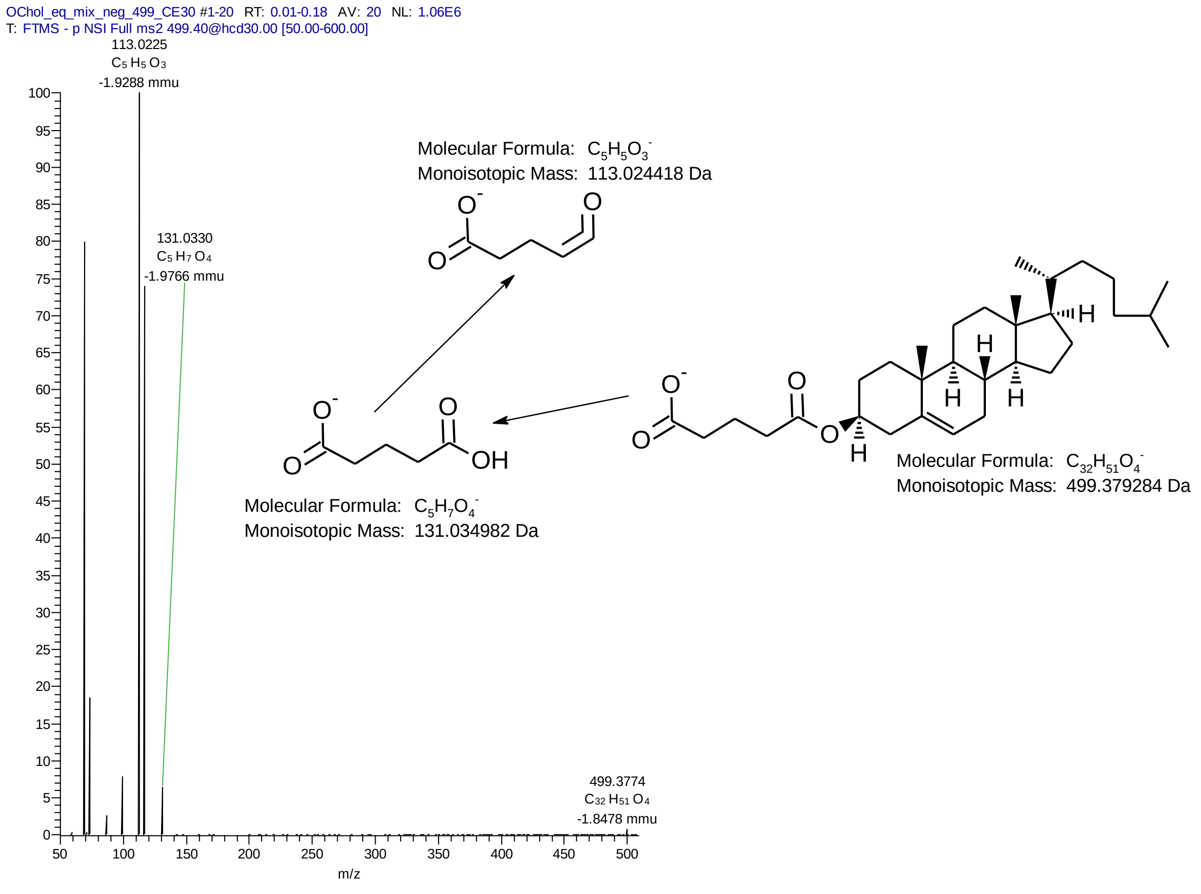

#### Supplementary Figure II Cont.

**C**

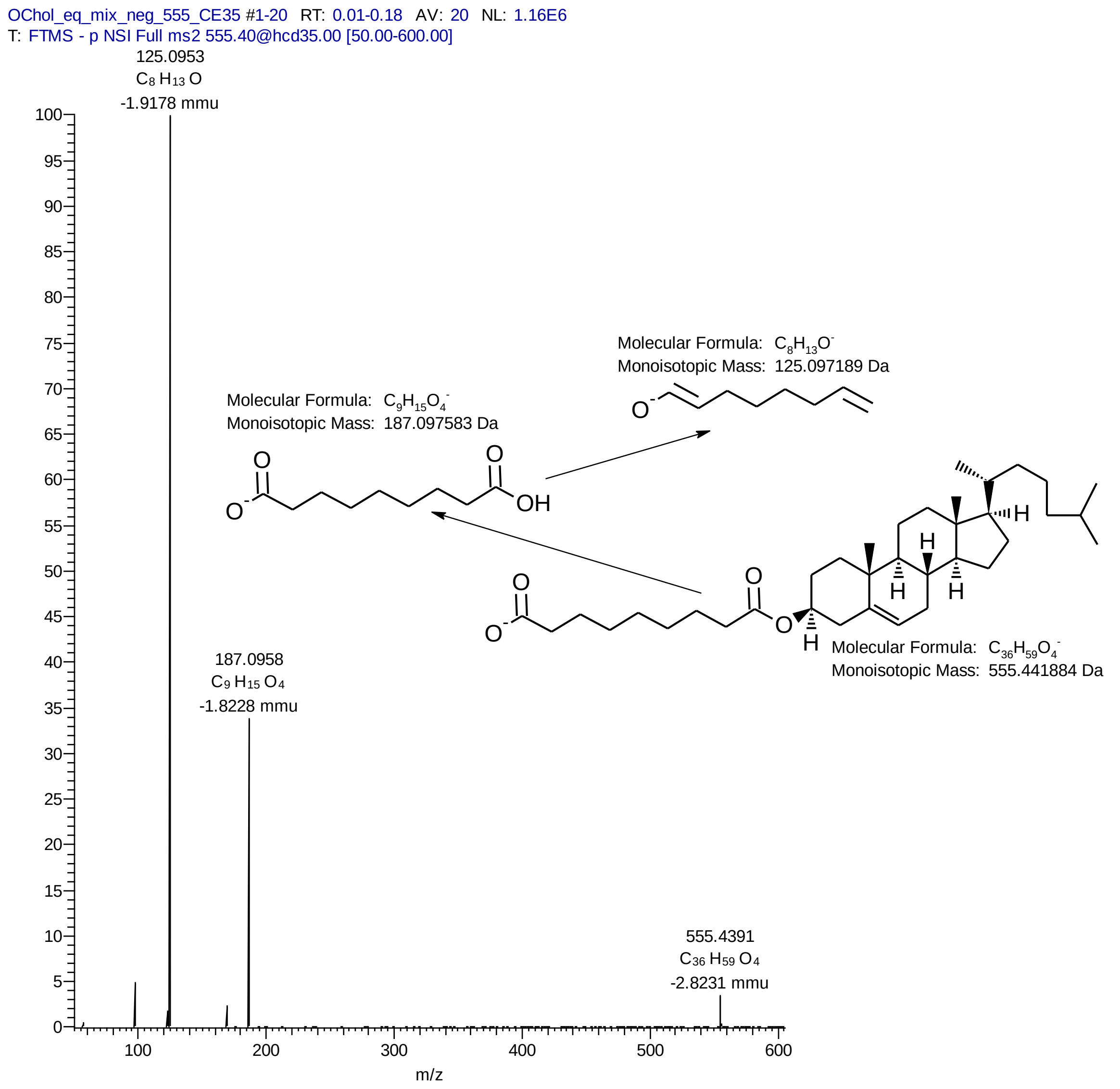


**Supplementary Figure II.**: MSMS spectra of **A.** ChS $[M-H^{+}]^{-}$ ion (at normalized ChE= 20%) and proposed fragmentation; **B.** ChS $[M-H^{+}]^{-}$ ion (at normalized CE= 30%) and proposed fragmentation, and **C.** ChA $[M-H^{+}]^{-}$ ion (at normalized ChE= 35%) and proposed fragmentation.

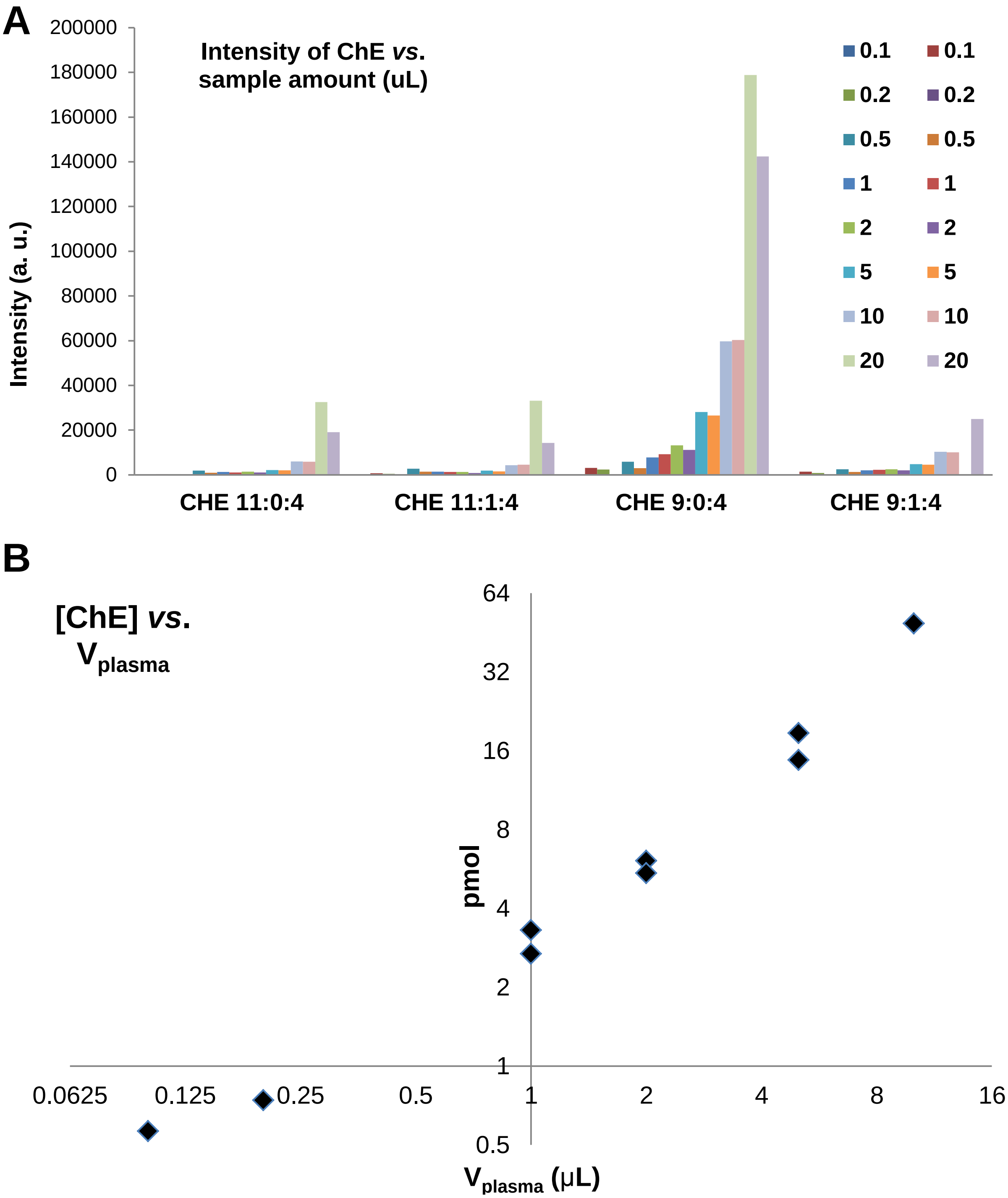


**Supplementary Figure III**. **A.** Different cholesteryl hemiesters (ChE) species observed in the test plasma sample and the dependence of their intensity with the volume of sample extracted (**B**).

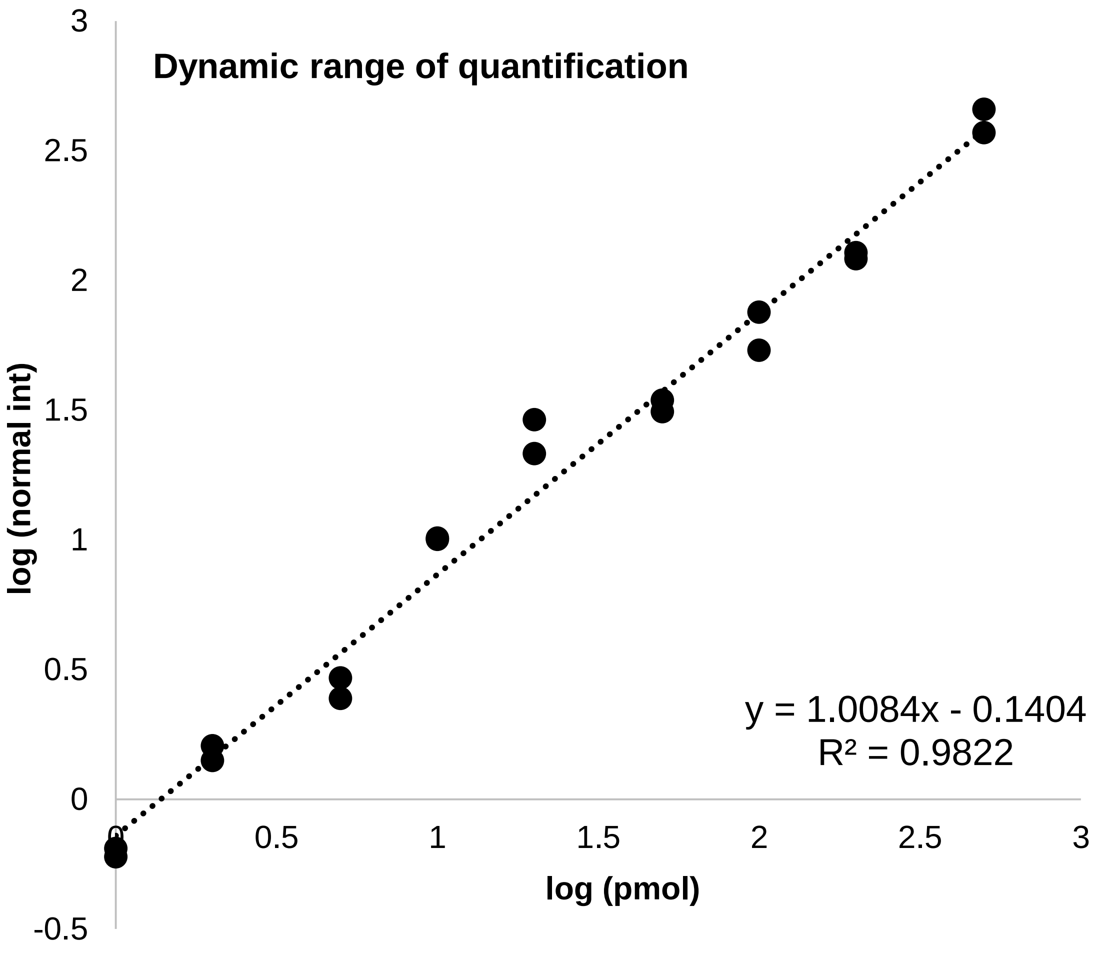


**Supplementary Figure IV.** ChE response with increasing amounts of standard added. The limit of quantification determined is 1 µM and the dynamic range is >500.

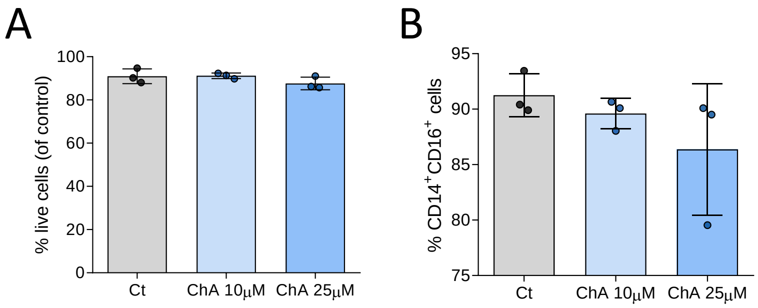


**Supplementary Figure V. ChA effect on viability and surface makers of monocytes (Mono) and monocyte-derived macrophages (MDM).** Peripheral blood was harvested from human volunteers. $CD14^+$ cells were purified from total mononuclear leukocytes (PBMCs) by flow cytometry. $CD14^+$ monocytes were cultured in the presence of ChA (10 μM or 25 μM) or vehicle (control) for 24 h (n=3) or for 7 days to obtain MDM (n=3). Monocytes were differentiated using rM-CSF. After lipid treatment, the percentage of live cells were determined (**A**) and cells were stained with monoclonal antibodies to determine surface expression of CD14 and CD16. **B**. Percentage of $CD14^+CD16^+$ positive cells after MDM differentiation (three different donors).

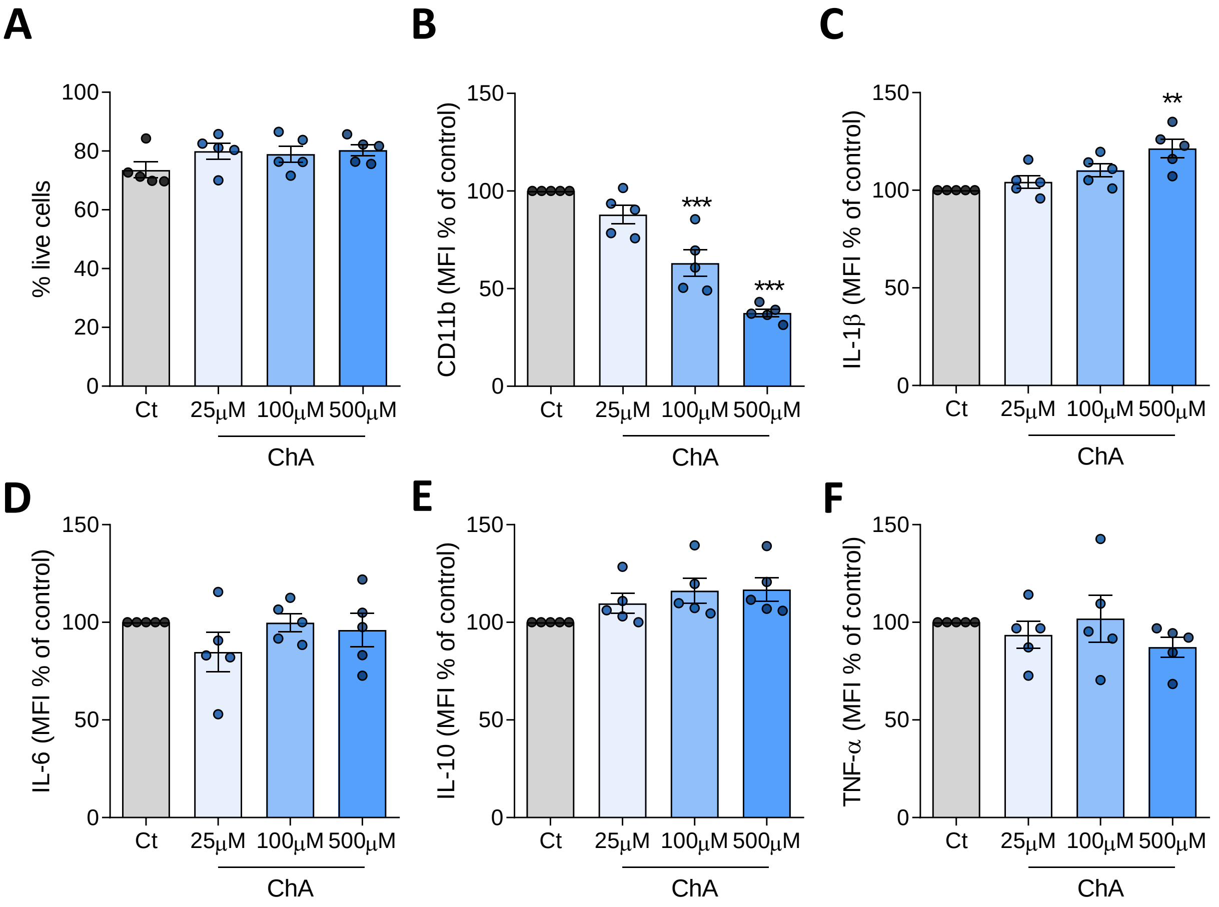


**Supplementary Figure VI. Impact of ChA on neutrophils viability and on the expression levels of inflammatory cytokines.** After isolation from the plasma of human healthy volunteers (n=5), neutrophils were cultured in the presence of ChA (25, 100 and 500 μM) or vehicle (POPC liposomes, control) for 4 h. Then, neutrophils were analyzed through flow cytometry to quantify the percentage of live cells (**A**) and the mean fluorescence intensity (MFI) ± SEM for CD11b (**B**) and inflammatory cytokines (**C**-**F**): IL-1β (**C**), TNF-α (**D**), IL-6 (**E**) and IL-10 (**F**). Data represent the mean ± SEM normalized to the control cells. **, *p* < 0.01, ***, *p* < 0.001.

### Supplementary Figure VII

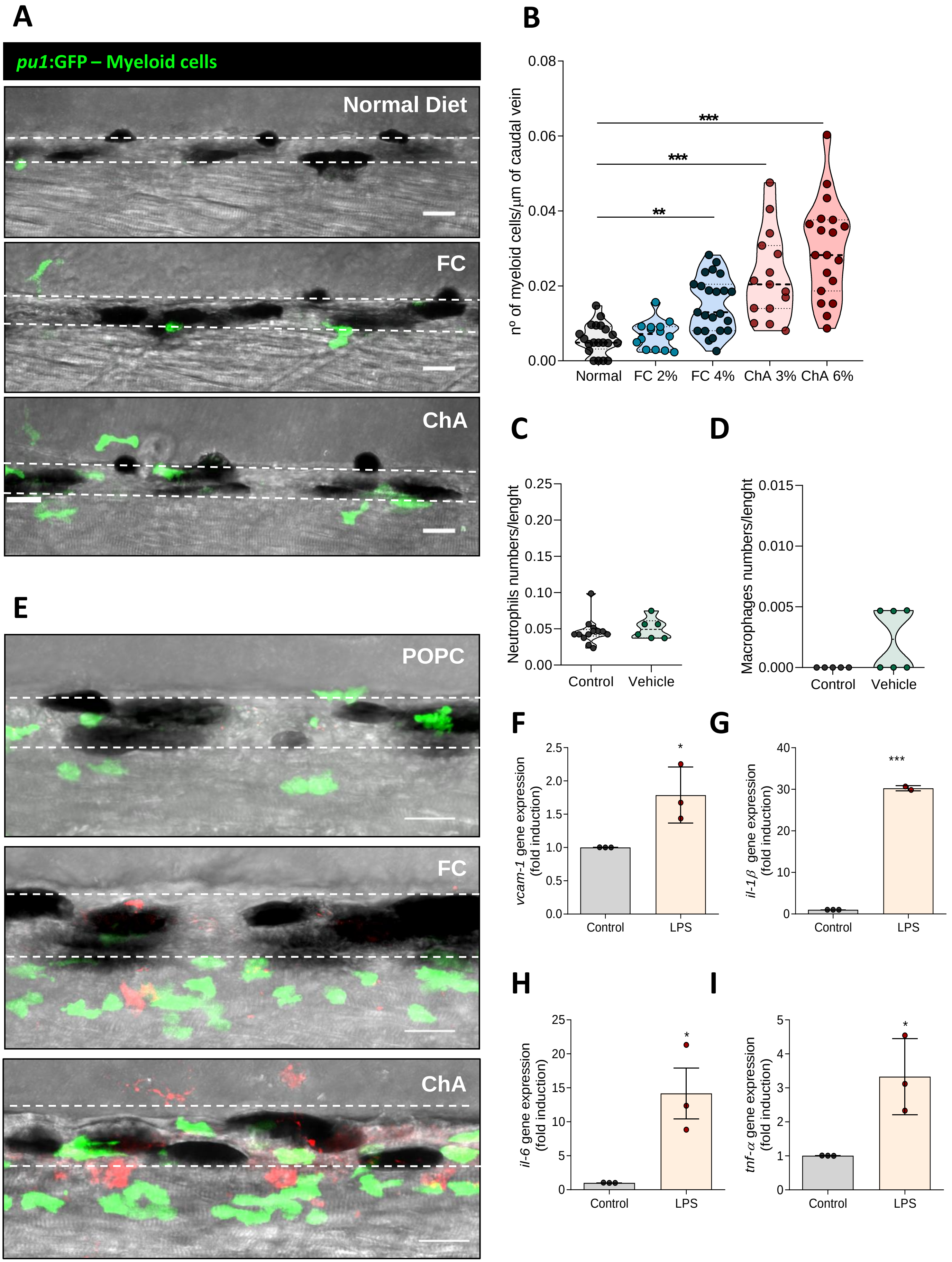

**Supplementary Figure VII. ChA induces myeloid cell infiltration into the vasculature.** Five days post-fertilization PU.1:EGFP larvae were fed with normal, FC- (2 and 4%) or ChA-enriched diet (3 and 6%) for 10 days. **A.** Merge of GFP-fluorescent cells (myeloid cells) and bright field images of the caudal vein (delineated by dotted lines). Scale bars, 20 µm. **B.** Quantification of myeloid cells normalized to the total lenght (µm) of caudal vein in larvae exposed to the different types of diet. **C, D.** Effect of embryonic medium (control) and POPC liposomes (27 µM) microinjection in infiltration of neutrophils and macrophages into the caudal vein. **E.** Representative images of neutrophils (green cells) and macrophages (red cells) recruitment into vasculature of Tg (mpeg.mCherryCAAX SH378, mpx:EGFP i114) larvae 24 h after lipid microinjection. **F-I**. Quantitative RT-PCR for *vcam-1* (**F**), il-1β (**G**), il-6 (**H**) ) and tnf-α (**I**) genes in zebrafish after LPS treatment for 24 h. The experiment was performed in triplicate (n = 20-30 zebrafish larvae per group) and the results are shown as mean ±SEM . $p^* < 0.05$; ** $p < 0.01$; ***, $p < 0.001$.

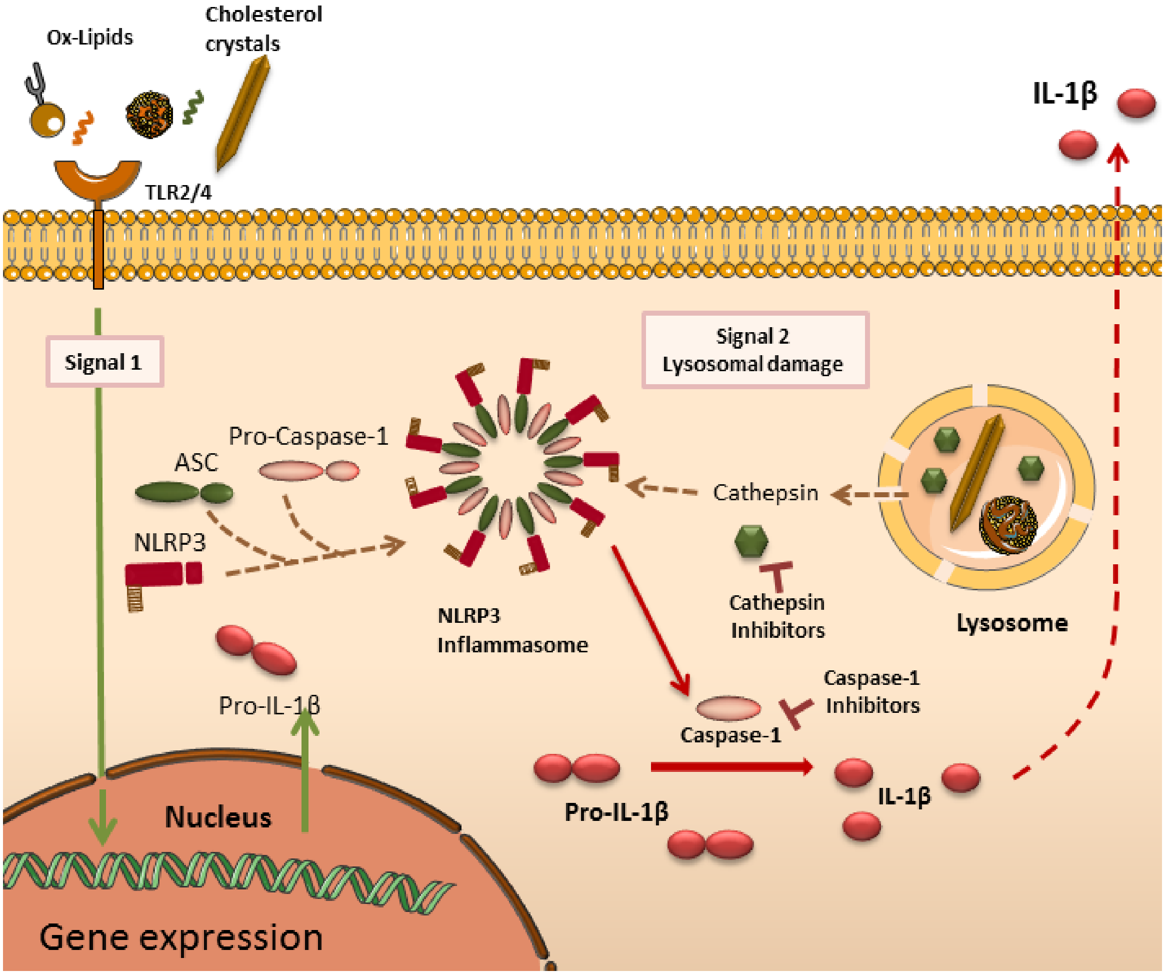


**Supplementary Figure VIII -** Schematic representation of inflammasome activation mechanism in context of atherosclerosis. To stimulate the release of IL-1β in macrophages, two distinct signals are necessary: first signal prime the cells (Signal 1) to activate the NF-κB-responsive genes *NLRP3* and *IL-1*β. In atherosclerosis, several lipids and lipoprotein agonists activate cell surface pattern recognition receptors (PRRs), such as Toll-like receptor 2 (TLR2) and TLR4, providing this signal 1. The second signal (Signal 2) promotes the assembly of the canonical NACHT, LRR and PYD domains-containing protein 3 (NLRP3) inflammasome complex. In atherosclerosis, Oxidized lipids, as ChA, can induce lysosomal membrane permeabilization, releasing lysosomal proteases, such as cathepsins. Cytosolic cathepsin activates NLRP3 with consequent maturation of caspase-1 which cleaves pro-IL-1β into bioactive IL-1β. IL-1β is then released from the cell. Several chemicals have been designed to inhibit the NLRP3 inflammasome, include C-YVAD-AOM. Cathepsin inhibitors (ZRLR) decrease IL-1β secretion by inhibiting NLRP3 inflammasome activation.

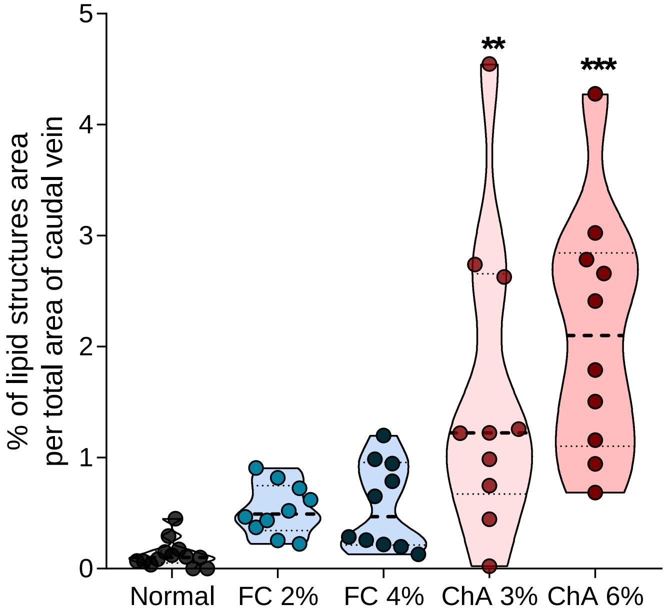


**Supplementary Figure IX: ChA induces accumulation of lipid in the vasculature earlier than free cholesterol.** After 5 days of feeding with normal, 2 or 4% FC-, 3 or 6% ChA-enriched diets, AB zebrafish larvae were imaged by confocal microscopy and the area of total lipid deposits area was quantified in zebrafish caudal vein. Fluorescent images of at least 10 larvae were quantified. The results are shown as mean ± SEM; **, $p < 0.01$; ***, $p < 0.001$.
